# Cryo-adapted bacterial copiotrophs from a Trans-Himalayan lake-desert ecosystem as biogeothermometers of warming, mitigators of perturbation, and candidates for biodegradation

**DOI:** 10.1101/2025.07.10.664119

**Authors:** Sumit Chatterjee, Subhajit Dutta, Jit Ghosh, Swapneel Saha, Mahamadul Mondal, Jagannath Sarkar, Nibendu Mondal, Wriddhiman Ghosh

**Affiliations:** Department of Biological Sciences, Bose Institute, Kolkata, India

**Keywords:** Trans-Himalayan deserts and lakes, psychrophilic and cryo-adapted bacteria, climate warming, antibiosis, microbiome protection, organic matter degradation

## Abstract

A Trans-Himalayan lake-desert ecosystem was explored for the low-to-high temperature adaptations of native copiotrophic psychrophiles that have the potentials for rendering substantive carbon remineralization under natural and/or anthropogenically influenced conditions of high organic matter delivery to the environment. Overall 27 species, classified across 15 genera of Actinomycetota, Bacillota, Bacteroidota, and Pseudomonadota, were isolated from the brackish-water and sediment-surface of Tso Moriri (a massive lacustrine body on the Changthang plateau that remains frozen for approximately one third of the year), and the fine talus covering a lake-side rocky mountain. In Luria broth, all the isolates grew at 4°C and 15°C; 13 of them grew, while 14 retained 19-99% cells in metabolically-active state, at −10°C; 23, 11, and none grew at 28°C, 37°C, and 42°C respectively. A *Cryobacterium* was most heat-susceptible, whereas some −10°C-growing *Arthrobacter*, *Microbacterium*, *Paenarthrobacter*, *Pseudomonas* and *Streptomyces* were most warming-resilient, among the present isolates. Catabolizing different simple/complex organic compounds, all isolates accomplished low/moderate/high growths at 4°C; 20 performed low growths at −10°C. Taxonomy/phylogeny did not determine the isolates’ growth/survival across the temperature-range tested. Lake-water and rock-dust isolates had the widest, while lake-sediment-dwellers had the narrowest, temperature-window for growth/survival. Rock-dust and lake-water isolates had more genes for low and high temperature adaptations, compared to the lake-sediment isolates. In co-cultures, most actinobacterial isolates inhibited the growth of mesophilic bacteria from warmer ecosystems, plus non-actinobacteria and actinobacteria from native/adjoining habitats. The data hold implications for biodegradation in cold/frigid territories, negative feedback control/reversal of warming, and microbiome protection from infiltration amidst climate change.

**IMPORTANCE:** Phylogenetically diverse, cold-adapted copiotrophic bacteria were retrieved from a Trans-Himalayan lake-desert ecosystem, and tested for chemoorganoheterotrophic growth at temperatures between −10°C and 42°C. Albeit growth dwindled at ≥28°C, most of the isolates thrived on simple as well as complex organic substances under freezing condition, which makes them prospective constituents of biodegradation reactors suitable for high-altitude/latitude deployment. In the natural habitats of the psychrophilic isolates, cessation of organotrophic activities at elevated temperatures would stop simple fatty acids and CO_2_ production, which in turn would preempt methanogenesis, undo greenhouse effect, and reverse environmental thawing (usher cooling). Such negative feedback control of homeostasis is fraught with the danger of microbiome takeover by thermally-better-adapted foreign organisms. Consequently, ecosystem preservation rests on the native microflora’s ability to resist external invasion. At 28°C, majority of the actinobacterial isolates inhibited bacteria from discrete higher-temperature ecosystems; they can, therefore, be viewed as the defenders of the cold/frigid ecosystem.

## INTRODUCTION

Microbial life at low temperatures is constrained by a number of biophysical and biochemical adversities (1, 2). However, across the seasonally or perpetually frozen alpine/polar territories of the Earth’s biosphere, barren soils, outwardly life-less rocky terrains, and most conspicuously, numerous limnic and fluvioglacial features, harbor considerably diversified microbiomes (3, 4) that in turn sustain highly climate-sensitive ecosystems via biogeochemical cycling of carbon and other elements (5–7).

Throughout the high-latitude and high-altitude areas of the globe, copious endolithic and chasmolithic microbial communities colonize the rocky terrains and contribute actively or passively to the weathering of rocks into dust and scree, thereby promoting sediment and soil formation (8, 9). Supraglacial and subglacial microbiomes drive considerable biogeochemical activities (10–12). The former types accelerate the melting of glaciers (13, 14), while post deglaciation the latter types release profuse greenhouse gases, such as carbon dioxide (CO_2_) and methane (CH_4_), into the atmosphere (15–17). Microbiomes thriving in young as well as matured soils, including those associated with permafrosts, and biocrusts covering deglaciated moraines and rock surfaces, render temporally slow but spatially extensive, fixation and remineralization of carbon (18, 19). Furthermore, approximately, 25 million lacustrine bodies (20), together with the vast expanses of thermokarst landforms, sequester huge amount of organic carbon by remaining frozen for a substantial part of the year, and emit enormous volumes of CO_2_ and CH_4_ upon thawing (21–25).

In all temporarily or permanently frozen ecosystems, microbial activities and proliferation apparently remain subdued as long as the temperature remains below or around 0°C (19, 26). However, organotrophic growth, and thereby remineralization of carbon and emission of greenhouse gases, starts once the temperature rises and cryoturbation takes place on a seasonal or climatic scale (19, 27–31). Greenhouse effect, at any spatiotemporal level, is thought to enhance microbial activities and growth, which in turn induces further thawing of the environment (19, 32). Such positive feedback cycles (27, 32, 33) can, in the long run, alter the structures and functions of microbiomes, and thereby entire ecosystems, within the cold/frigid realm.

To understand the functioning of cold/frigid ecosystems, and in order to manage them in a sustainable way, we require comprehensive information on the scopes of carbon cycling and sequestration around the freezing point of water. For that purpose, chemoorganoheterotrophic capabilities of autochthonous microorganisms need to be delineated at near-zero and sub-zero degree Celsius, via extensive pure-culture-based investigations. Furthermore, in the context of global warming in the present era, it is imperative to appreciate the homeostatic vulnerabilities of alpine/polar microbiomes alongside their potential intrinsic resilience against perturbations. This would subsequently help mitigate impending microbiome (and habitat) alterations triggered by climate change across the global cryosphere. Towards that end, we require comprehensive data on how psychrophilic microorganisms (34–36) respond to different levels of warming in terms of population-level growth or survival. Concurrently, in relation to plausible microbiome transformation within cold/frigid ecosystems due to global warming, we need to know whether cold-adapted microbial communities have any indigenous safeguard against infiltration of organisms from warmer territories (37, 38).

The present study of pure-culture microbiology explored three adjacent environments (habitats) within a Trans-Himalayan lake-desert ecosystem, centered on the massive fresh-to-brackish water body called Tso Moriri (vernacular meaning: lake amid the mountains), which is situated on the cold arid Changthang plateau of eastern Ladakh, India (Figs. 1a-c). Phylogenetically diverse chemoorganoheterotrophic bacterial psychrophiles were isolated and characterized from Tso Moriri’s sediment, and water that remains frozen for approximately one third of the year. Psychrophilic heterotrophs were also isolated from the weathered rock dust (fine talus and scree particles) that covers the hill overlooking the western bank of the lake (Figs. 1d-g). Prior to the isolation of pure cultures, the water, sediment and talus samples were subjected to iterative cycles of freezing and thawing. This ensured that the bacterial strains retrieved were cryo-adapted (capable of rendering growth, whether little or substantial, at −10°C), or at least cryo-tolerant (adept in population-level survival, i.e. retention of >1% cells in metabolically-active state, at −10°C), in nature.

**Fig. 1.**
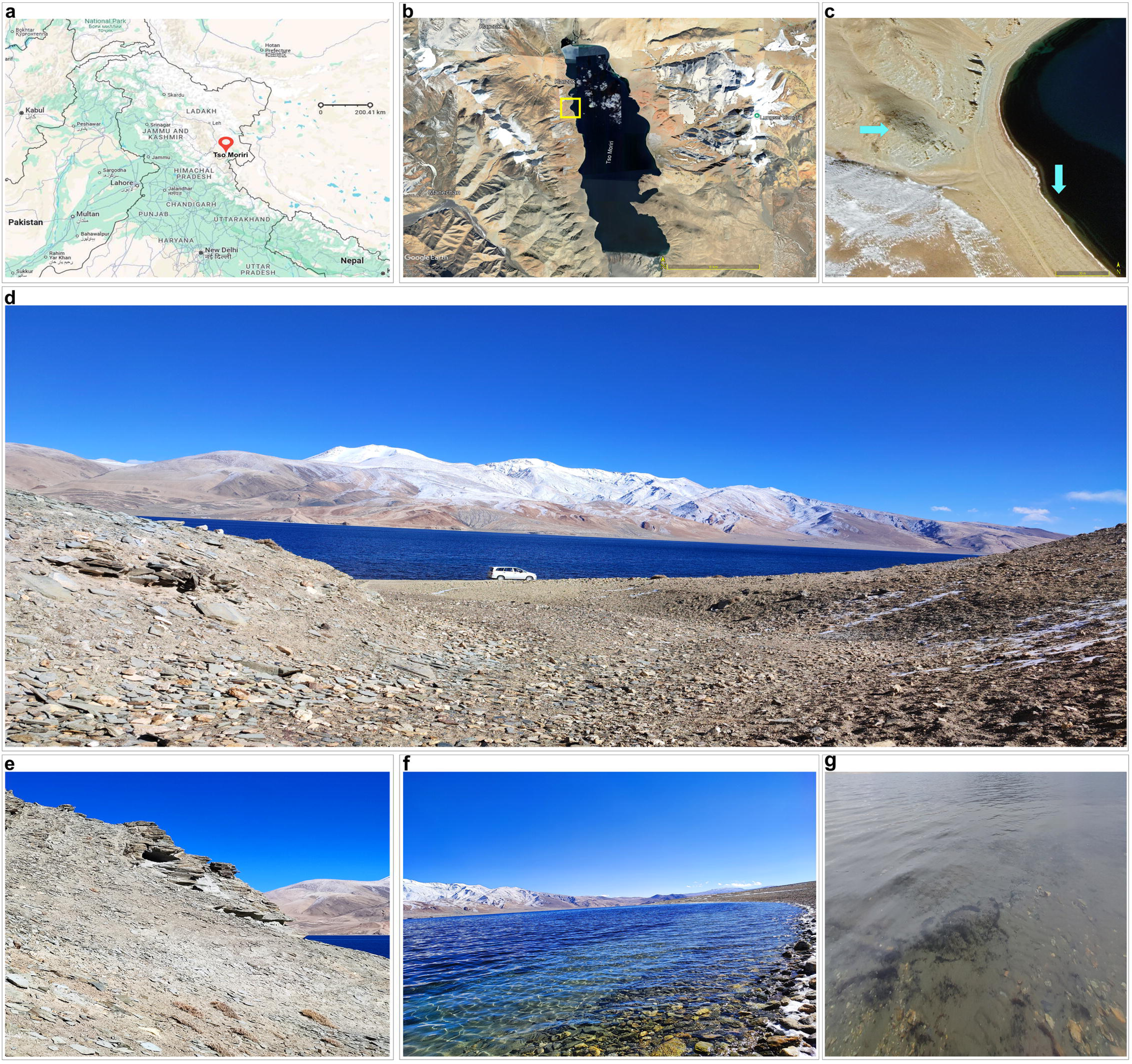
Geographical location of the Tso Moriri lake, topography of the explored area, and position of the sampling sites. (**a**) Map of northern India showing the location of the Tso Moriri lake. (**b**) Satellite view of the lake shown in the context of the adjoining areas of the Changthang desert (position of the study area is indicated by a yellow square). (**c**) Satellite view of the west bank of Tso Moriri where the study sites (indicated by cyan arrows) were situated. (**d**) Panoramic view of the portion of the lake and the adjoining hill from where samples were collected. (**e**) The barren hill slope where weathered rock dust (fine talus and scree) was sampled for the upper 1 cm from the surface. (**f**) Shallow coastal water of Tso Moriri that was sampled from within a depth of 1 cm. (**g**) Shallow coastal sediment of Tso Moriri that were sampled for the top 1 cm. Maps and views shown in panels (**a**), (**b**) and (**c**) were sourced from the open database located at https://www.google.co.in/maps.

A separate time-series investigation of biogeochemistry, carried out by our laboratory (Chatterjee et al., unpublished) at the same sample sites explored in this study, had shown that after the summer months (just before the winter), concentration of total carbon in the surficial lake-water reached 80 mg L^-1^, while total carbon content of the surficial samples of lake-sediment and weathered rock dust reached approximately 2% (w/w) and 1% (w/w) respectively. In view of these findings [notably, the aforesaid numbers are reasonably close to the carbon content values reported from many tropical marine ecosystems (39, 40)], the present enrichment and isolation strategy specifically targeted cryo-adapted, or cryo-tolerant, copiotrophs that possess the potentials for rendering substantive carbon remineralization under cold and frigid conditions. The overall objective was to ensure that the organotrophic growth potentials exhibited by the new isolates *in vitro* held direct implications for the scope of *in situ* organic matter degradation across alpine/polar ecosystems experiencing high carbon delivery to the environment, naturally and/or under anthropogenic influence.

In the aforesaid context, it was first investigated whether the copiotrophic psychrophiles isolated in nutrient-rich medium could grow by catabolizing different simple or complex organic substrates at near-zero and sub-zero degree Celsius. Experiments were then conducted to know how the growth and activity of the isolates were affected by different levels of warming. The data obtained *in vitro* were evaluated theoretically to explore whether natural populations of the isolated bacteria could act as “biogeothermometers” chronicling *in situ* temperatures over time and space. From the contemporary perspective of heightened thawing of the cold/frigid realm, we examined whether the bacteria retrieved from the Tso Moriri area (TMA) had any aptitude for resisting incursion of foreign microorganisms from warmer climatic territories. For this purpose we tested the antibiosis potentials of the current isolates against higher-temperature-adapted bacteria retrieved previously from discrete warmer ecosystems. All phenotypic features delineated for the new isolates were appraised in the light of the organisms’ genome contents. The whole repertoire of data available was eventually interpreted in the context of climate change to envisage if the indigenous psychrophiles had any role in the abatement of environmental warming. Cold/frigid ecosystems, in the present era, being exceedingly perturbed by anthropogenic waste deposition (41) possibilities were weighed for using the copiotrophic TMA psychrophiles in bioreactors designed to degrade organic litters at near-zero and sub-zero degree Celsius.

## RESULTS

### Cold-adapted copiotrophs from Tso Moriri lake-desert ecosystem

A sum total of 61 aerobic, copiotrophic, and freeze-thaw resilient bacterial pure-cultures, apparently dissimilar with regard to colony morphology, were enriched and isolated in Luria broth (LB), from the lake-water (Fig. 1f), lake-sediment (Fig. 1g), and rock-dust (Fig. 1e) samples. 22 of these were obtained from the weathered rock dust of the lake-side hill, while five and 34 were from the lake’s water and sediment respectively (Table 1). 16S rRNA gene sequence similarities clustered the 61 isolates into 27 species-level entities that in turn were ascribable to 15 genera distributed over the phyla Actinomycetota, Bacillota, Bacteroidota and Pseudomonadota. One representative strain from each species-level cluster was selected for further characterization (Table 1).

**Table 1.**
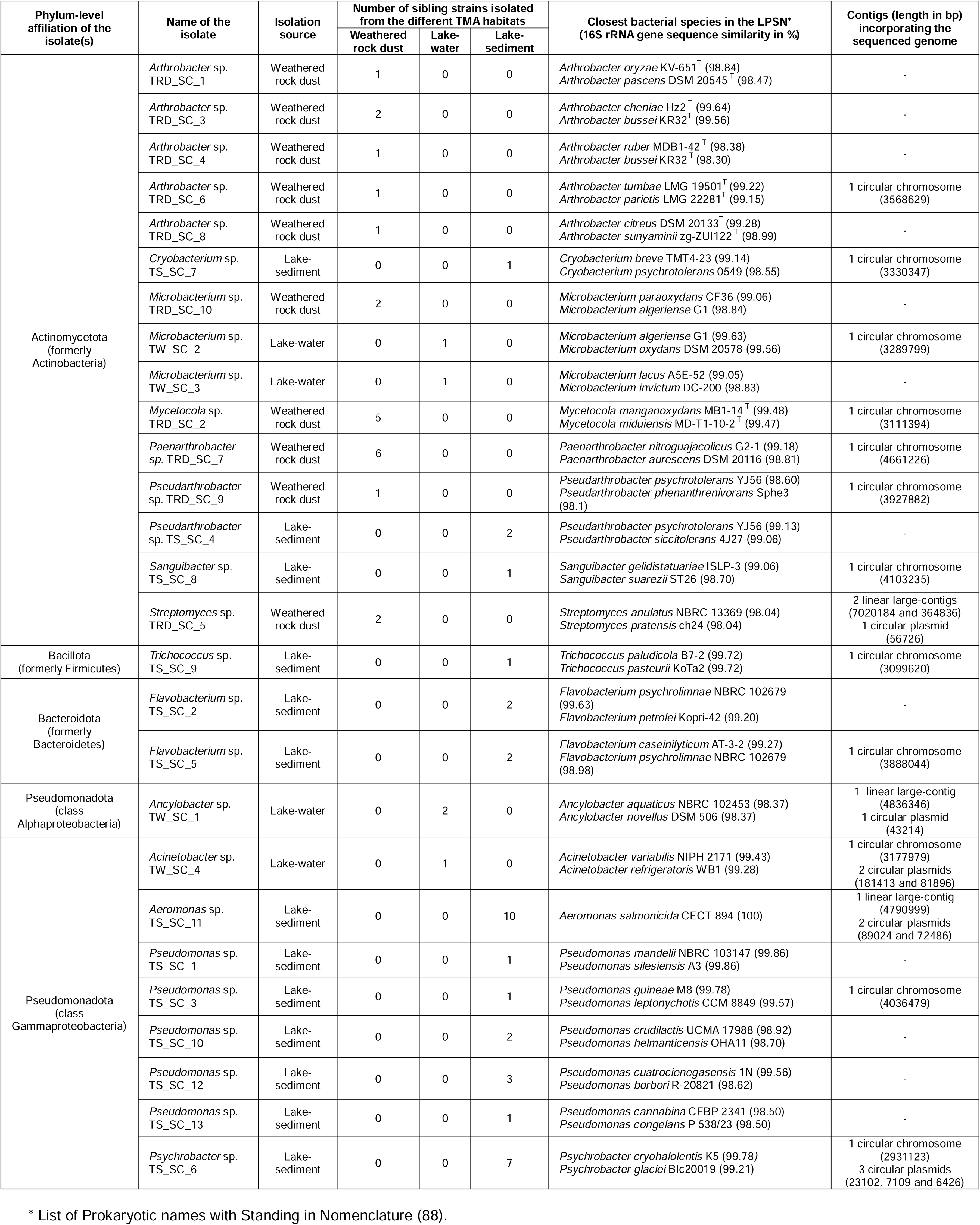
Chemoorganoheterotrophic bacteria isolated from three different environments (habitats) within the Tso Moriri area (TMA), viz. lake-water, lake-sediment, and weathered rock dust of a lake-side hill.

### Cold adaptations of the TMA isolates

In LB medium, all the 27 species retrieved from the Tso Moriri area could grow at 4°C (Figs. 2c-d) and 15°C (Figs. 2e-f), whereas only 13 could grow at −10°C (Figs. 2a-b). In terms of what percentage of the starting CFU density (colony-forming units mL^-1^) remained in the culture after the stipulated period of incubation (Tables S1-S3), the extents of growth recorded at −10°C were far lower than those recorded at 4°C and 15°C. After 14 days at 4°C, the cultures of *Flavobacterium* TS_SC_2 and *Arthrobacter* TRD_SC_3 showed the lowest and highest increases in CFU density respectively. After two days at 15°C, lowest and highest increases in CFU density were exhibited by the cultures of *Flavobacterium* TS_SC_2 and *Arthrobacter* TRD_SC_8 respectively. After 28 days at −10°C, *Pseudarthrobacter* TRD_SC_9 and *Arthrobacter* TRD_SC_6 had the lowest and highest increases in CFU density respectively.

**Fig. 2.**
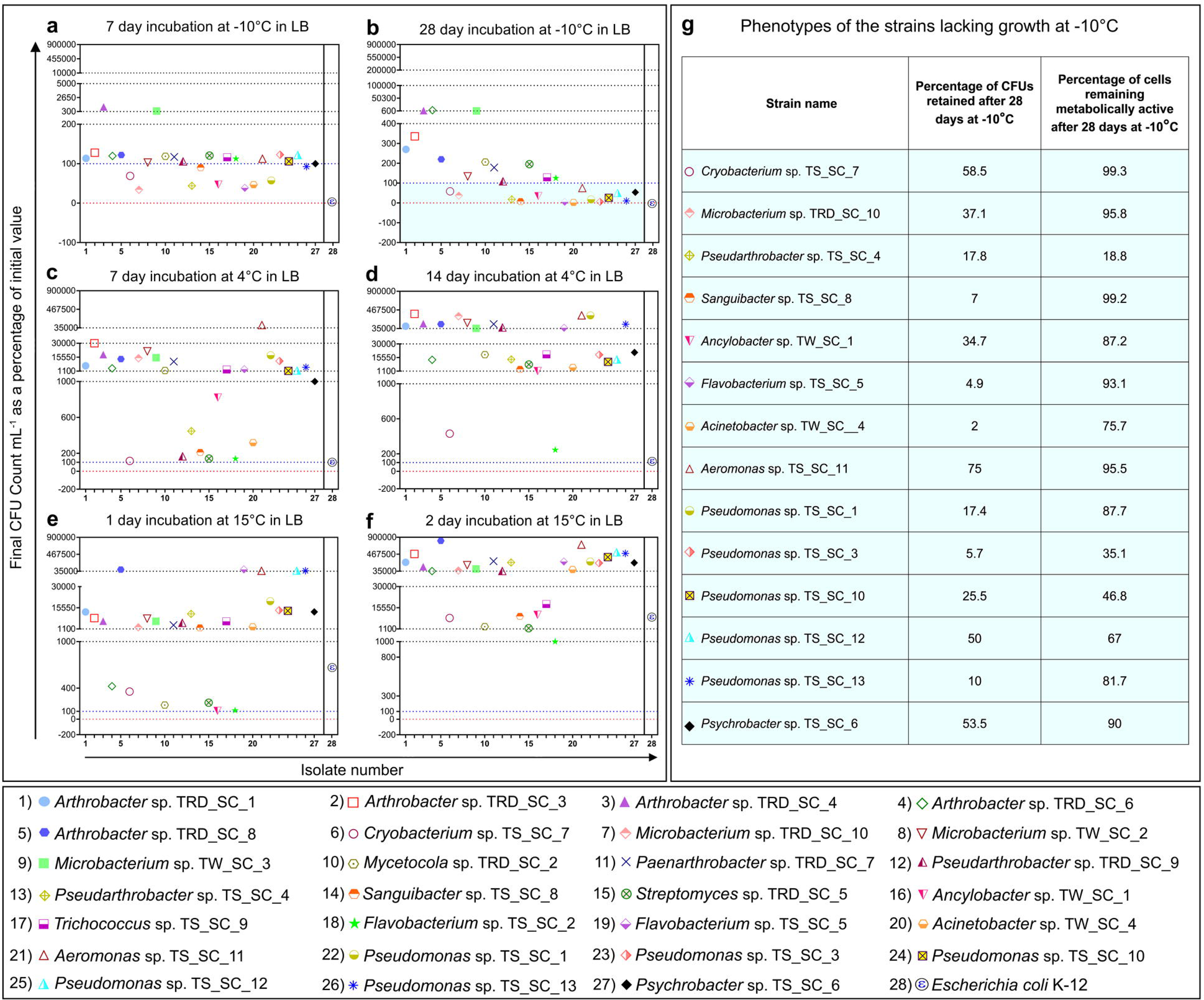
Growth, survival, or abolition of cell populations documented for the TMA isolates in LB medium under cold or frigid conditions. Growth, or the lack of it, was tested at (**a** and **b**) −10°C, (**c** and **d**) 4°C, or (**e** and **f**) 15°C, and expressed as the increase or decrease in CFU density recorded over time (final CFU count mL^-1^ of the culture as a percentage of the initial value). Every number plotted is an arithmetic mean of the data acquired from three different experiments, where standard deviation always remained <2% of the mean. Numerical values of the CFU densities underlying the data shown in panels (**a** and **b**), (**c** and **d**), and (**e** and **f**) can be found in Tables S1, S2, and S3 respectively. (**g**) Percentage of the initial CFU density retained post incubation by each isolate lacking growth at −10°C is tabulated alongside the percentage of cells remaining metabolically active (capable of taking FDA stain) within the same culture. Refer to Fig. S1 for the flow cytometry-derived dot plots depicting what proportions of cells were stained by FDA in the 14 different isolates incapable of growing at −10°C.

Notably, the 14 species, which could not grow in LB at −10°C, managed to maintain considerable proportions of their cell populations in divisible (Figs. 2a-b) and/or metabolically-active (Figs. 2g and S1) states, as evidenced by CFU density data, and fluorescein diacetate (FDA) staining followed by flow cytometry, respectively. After 28 days at this temperature, *Acinetobacter* TW_SC_4 and *Aeromonas* TS_SC_11 retained the lowest and highest proportions (2% and 75%) of the initial CFU density respectively, while *Pseudarthrobacter* TS_SC_4 had the lowest (19%), and *Cryobacterium* TS_SC_7 and *Sanguibacter* TS_SC_8 had the highest (99%), proportions of cell stained with FDA. Furthermore, for every bacterium lacking growth at −10°C, the percentage of metabolically-active cells exceeded the percentage of cells retaining their divisibility after 28 days of incubation, even though no significant correlation existed between the proportions of divisible and metabolically-active cells (Fig. S2a).

### Growth on different simple/complex carbon compounds at zero and sub-zero degree Celsius

After 14 days of incubation at 4°C, all the bacterial species isolated from the TMA rendered at least a little growth by utilizing no less than three out of the 10, simple or complex, organic compounds tested as single chemoorganoheterotrophic substrates (Tables S4-S13). The extent of growth differed across the isolates in terms of the percentage increase in CFU density that was attributable to the utilization of the carbon compound in question (Fig. 3a; Table S14). 10 TMA isolates were found to accomplish low, moderate, or high growth on all the 10 compounds tested; these were *Aeromonas* TS_SC_11, *Arthrobacter* TRD_SC_3, *Cryobacterium* TS_SC_7, *Flavobacterium* TS_SC_2, *Flavobacterium* TS_SC_5, *Microbacterium* TRD_SC_10, *Sanguibacter* TS_SC_8, *Pseudomonas* TS_SC_12, *Pseudomonas* TS_SC_13 and *Psychrobacter* TS_SC_6. Although unable to use acetate, *Streptomyces* TRD_SC_5 could accomplish high growth on maximum number of complex carbon compounds, namely cellulose, chitin, pectin, starch and xylan; this organism also rendered moderate growth on agar, albumin and hexadecane, while its growth on benzoate was low. In contrast, *Pseudomonas* TS_SC_3, which could only render moderate growth on cellulose, and low growth on albumin and starch, was the least efficient TMA isolate in terms of chemoorganoheterotrophic utilization of the carbon substrates tested.

**Fig. 3.**
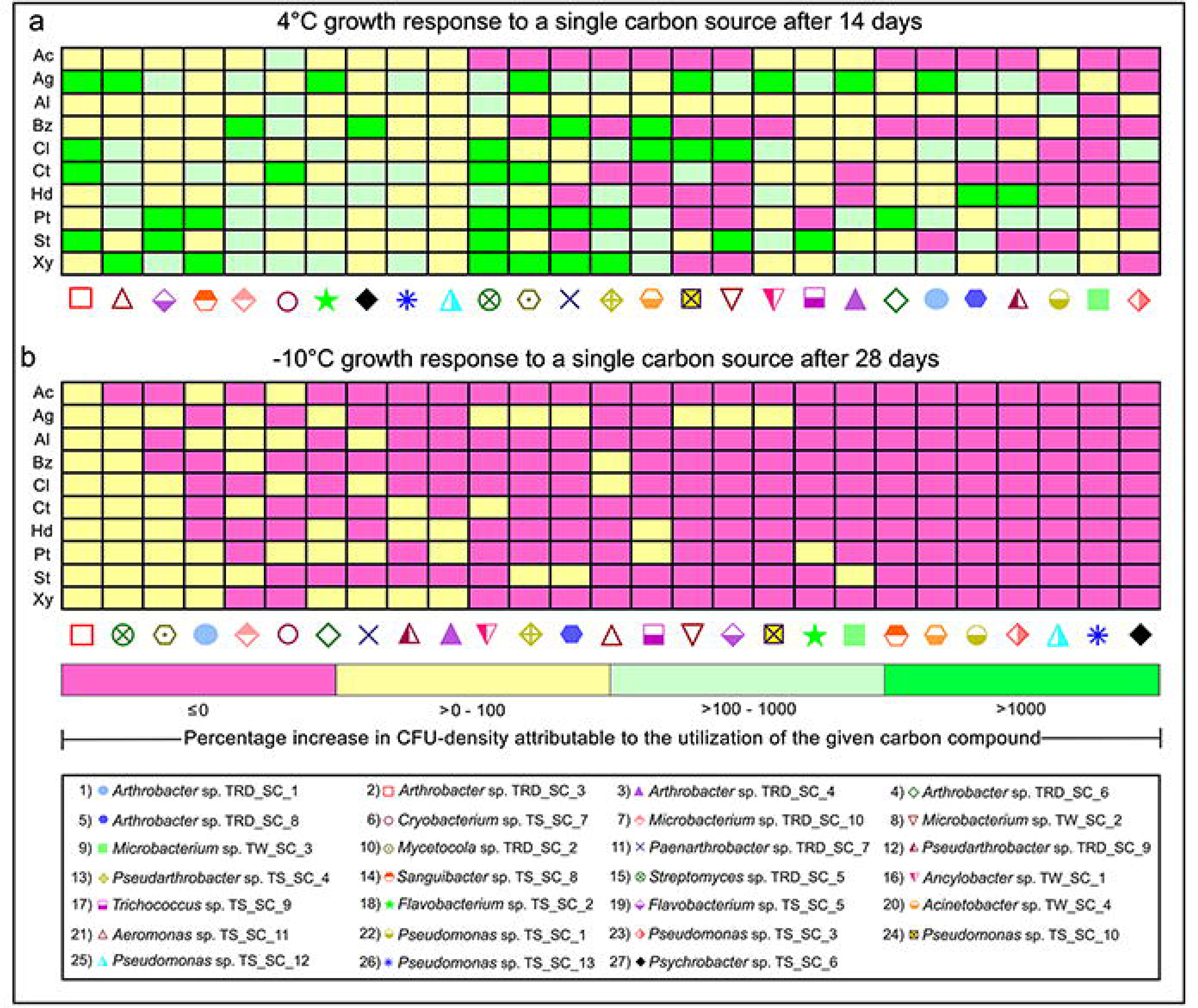
Abilities of the 27 TMA isolates to grow at near-zero and sub-zero degree Celsius temperatures in MS solution supplemented with one simple or complex carbon compound at a time: **Ac**, acetate; **Ag**, agar; **Al**, albumin; **Bz**, benzoate; **Cl**, cellulose; **Ct**, chitin; **Hd**, hexadecane; **Pt**, pectin; **St**, starch; **Xy**, xylan. After incubation for (**a**) 14 days at 4°C, or (**b**) 28 days at −10°C, growth was considered to have occurred due to the utilization of a given carbon compound only when a positive remainder value was obtained after subtracting the percentage increase in CFU density recorded for the same organism in MS (here also, only positive values were considered) from the percentage increase in CFU density recorded in MS supplemented with the carbon compound in question (here too, only positive values were considered). Numerical values for the percentage increases in CFU-density that were attributable to the utilization of the different carbon compounds at 4°C and −10°C have been given in Tables S14 and S25, while all CFU density values underlying the data shown in Tables S14 and S25 have been presented in Tables S4-S13 and S15-S24, respectively.

After 28 days of incubation at −10°C, a sum total of 20 TMA isolates were found to render low but definite growth on at least one of the 10 carbon compounds tested (Tables S15-S24), even though in LB medium, only 13 species had exhibited low to high growth at −10°C (Figs. 2a-b; Table S1). Of the 14 TMA isolates that had failed to render −10°C-growth in LB, seven – namely, *Aeromonas* TS_SC_11 (benzoate and cellulose), *Ancylobacter* TW_SC_1 (agar and chitin), *Cryobacterium* TS_SC_7 (acetate, albumin, cellulose and pectin), *Flavobacterium* TS_SC_5 (agar), *Microbacterium* TRD_SC_10 (agar, albumin, benzoate, chitin and starch), *Pseudarthrobacter* TS_SC_4 (agar and starch), and *Pseudomonas* TS_SC_10 (agar) – could render a low level of growth attributable to the use of at least one carbon compound (Fig. 3b; Table S25). Overall, *Arthrobacter* TRD_SC_3 accomplished low levels of growth on all the 10 compounds tested, while *Streptomyces* TRD_SC_5 rendered low levels of growth on all the substrates except acetate, and *Mycetocola* TRD_SC_2 could do the same on all the carbon compounds except acetate, albumin and benzoate. Of the remaining 24 isolates, 17 could achieve low levels of growth on only a few, or just one, of the 10 organic substrates tested; seven isolates could grow on none of them (Fig. 3b; Table S25).

### Extreme oligotrophy at zero and sub-zero degree Celsius

In the above experiments, growth of an isolate on a given organic substrate was defined exclusively by a positive remainder value obtained after subtracting the percentage increase in CFU density recorded for the isolate in MS solution from the percentage increase in CFU density recorded in MS plus the substrate in question (furthermore, for the two percentage terms in question, only positive values were taken into consideration). In that context, a separate scrutiny of the increases in the CFU densities of the different isolates recorded after incubation in MS solution at 4°C and −10°C revealed their extreme oligotrophic potentials.

At 4°C, after 14 days of incubation in MS solution (Table S26), CFU densities of *Microbacterium* TW_SC_2, and *Pseudomonas* strains TS_SC_1 and TS_SC_10, increased by >200% of the initial levels, whereas those of *Acinetobacter* TW_SC_4, *Arthrobacter* TRD_SC_1 and TRD_SC_6, *Microbacterium* TW_SC_3, *Mycetocola* TRD_SC_2, *Paenarthrobacter* TRD_SC_7 and *Trichococcus* TS_SC_9 increased by 6-72%. Under these conditions, CFU densities of *Arthrobacter* TRD_SC_3, *Microbacterium* TRD_SC_10, and *Streptomyces* TRD_SC_5 remained unchanged, while those of the remaining 14 strains decreased by 3-90% of the initial levels.

At −10°C, after 28 days in MS (Table S27), CFU densities of *Arthrobacter* strains TRD_SC_1 and TRD_SC_6, *Microbacterium* strains TW_SC_2 and TW_SC_3, *Mycetocola* TRD_SC_2, and *Paenarthrobacter* TRD_SC_7 increased by 10-50%, whereas those of the remaining species decreased by 2-98%, of the corresponding initial levels.

### Progressive thermal susceptibility of the TMA isolates

In LB medium, 23 out of the 27 TMA isolates studied could grow at 28°C (Figs. 4a-b; Table S28), whereas only 11 could grow at 37°C (Figs. 4c-d; Table S29), and none at 42°C (Figs. 4e-f; Table S30). In terms of what percentage of the starting CFU density remained in the culture after the stipulated period of incubation, extents of growth recorded at 28°C were much higher than those recorded at 37°C.

**Fig. 4.**
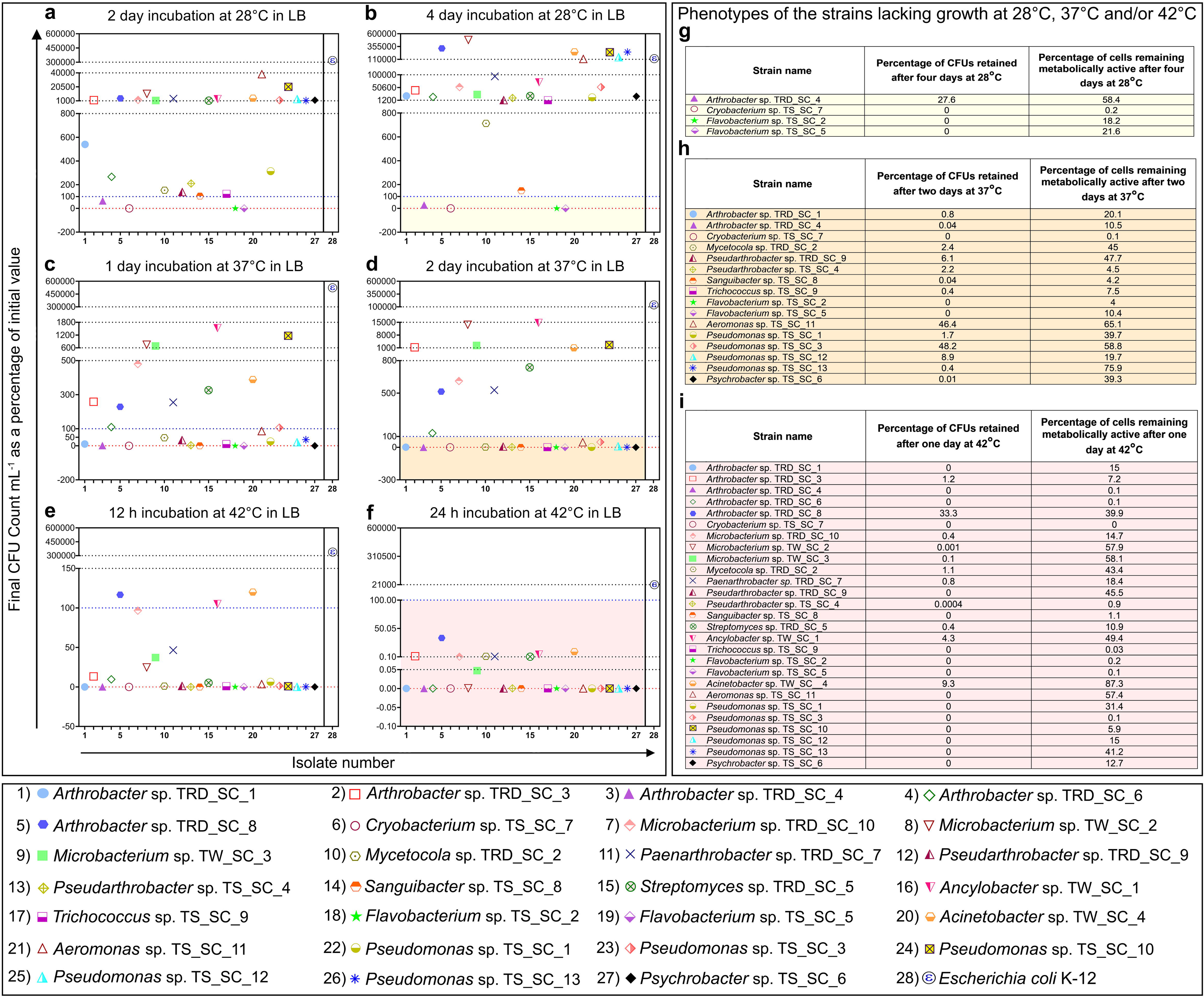
Growth, survival, or abolition of cell populations documented for the TMA isolates in LB medium at temperatures above the psychrophilic range. Growth, or the lack of it, was tested at (**a** and **b**) 28°C, (**c** and **d**) 37°C, or (**e** and **f**) 42°C, and expressed as the increase or decrease in CFU density recorded over time (final CFU count mL^-1^ of the culture as a percentage of the initial value). Every number plotted is an arithmetic mean of the data acquired from three different experiments, where standard deviation always remained <2% of the mean. Numerical values of the CFU densities underlying the data shown in panels (**a** and **b**), (**c** and **d**), and (**e** and **f**) can be found in Tables S28, S29, and S30 respectively. (**g**, **h**, and **i**) Percentages of the initial CFU densities retained post incubation by each isolate lacking growth at 28°C, 37°C, and 42°C respectively are tabulated alongside the percentages of cells remaining metabolically active (capable of taking FDA stain) within the same cultures. Refer to Figs. S3a, S3b, and S4 for the flow cytometry-derived dot plots depicting what proportions of cells were stained by FDA in the 4, 16, and 27 different isolates incapable of growing at 28°C, 37°C, and 42°C respectively.

Of the four species incapable of growing in LB at 28°C, three did not retain any CFU, while only *Arthrobacter* TRD_SC_4 retained 28% of the initial CFU density, after four days of incubation at this temperature. At the same time, three of these four isolates had 18-58% cells in metabolically-active conditions, while *Cryobacterium* TS_SC_7 had 0.2% cells in active condition (Figs. 4g and S3a).

At 37°C, 16 TMA isolates failed to grow in LB. Nine of them had zero or near-zero CFU density, while the other seven had 1-48% of the initial CFU densities, remaining in the cultures after two days of incubation. At the same time, 15 out of these 16 isolates had 4-76% cells in metabolically-active conditions, while *Cryobacterium* TS_SC_7 had 0.1% cells in active condition (Figs. 4h and S3b).

None of the 27 species retrieved from the Tso Moriri area could grow in LB at 42°C. 22 of them had zero or near-zero CFU density, while the remaining five had 1-33% of the initial CFU densities, present in the cultures after one day of incubation. At the same time, 19 out of the 27 isolates had 1-87.3% cells in metabolically-active conditions, while eight had <1% cell in active condition (Figs. 4i and S4).

Overall, the trends of LB-based population-level survival of the isolates lacking growth at ≥28°C showed that the percentage of cells remaining metabolically active at a given high temperature exceeded the percentage of cells which retained their divisibility at that temperature (Figs. 4g-i). Furthermore, for all strains susceptible to high temperatures, proportions of divisible, and metabolically active, cells remaining in the cultures after stipulated periods of incubation decreased with increase in temperature. That said, no significant correlation existed between the proportions of divisible and metabolically-active cells, across the species incapable of growth at high temperatures (Figs. S2b-d).

### Differential temperature windows for growth and survival of cell populations

At every incubation temperature tested for growth in LB medium (−10°C, 4°C, 15°C, 28°C, 37°C, or 42°C), different TMA isolates exhibited different rates of increase or decrease in CFU density (Tables S31-36 show the data for the six individual temperatures). On the flip side, temperature windows for growth (increase in CFU density) and population-level survival (retention of >1% cells in metabolically-active state), over the tested range, varied across the isolates (Fig. 5). *Arthrobacter* TRD_SC_3 and TRD_SC_8, *Microbacterium* TW_SC_2 and TW_SC_3, *Paenarthrobacter* TRD_SC_7, and Streptomyces TRD_SC_5, shared the widest temperature range over which growth was recorded (−10°C to 37°C), whereas *Flavobacterium* TS_SC_5 and *Cryobacterium* TS_SC_7 had the narrowest growth window (4°C to 15°C).

**Fig. 5.**
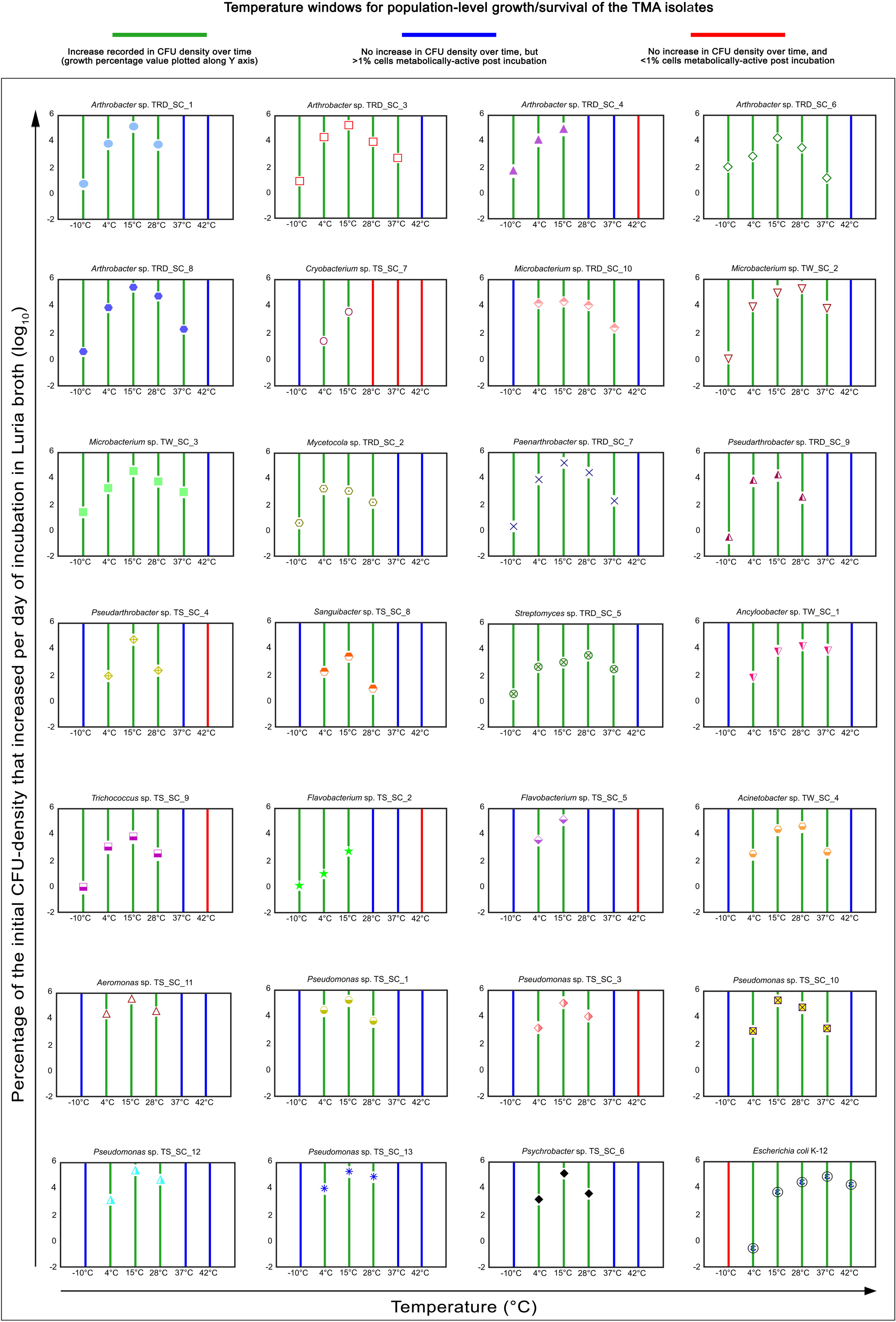
Growth rates (percentages of the initial CFU densities that increased per day of incubation), or absence of growth (no increase in the CFU density over time), recorded for the individual TMA isolates in Luria broth at −10°C, 4°C, 15°C, 28°C, 37°C, and 42°C. Within a given panel, a green bar indicates an increase recorded in the CFU density over time (growth). Blue bar indicates no increase in CFU density over time, but >1% cells of the culture getting stained with FDA at the end of the incubation. Red bar indicates no increase in CFU density over time, and <1% cells of the culture getting stained with FDA at the end of the incubation. Numerical values for all the CFU densities underlying these graphical representations can be found in Tables S31-S36. FACS data for the proportions of FDA-stained cells can be found in Figs. S1, S3 and S4.

For all those 4°C-growing isolates whose growth in LB ceased at −10°C, more than 1% cells remained metabolically active at this freezing temperature (Fig. 2g); the −10°C-growing isolates, on the other hand, were likely to remain metabolically active at even lower temperatures. At ≥28°C, all the TMA isolates, except *Arthrobacter* TRD_SC_6 and *Cryobacterium* TS_SC_7, exhibited population-level survival at temperatures above the points where their growth was last recorded (Figs. 5 and 6a). For TRD_SC_6 and TS_SC_7, growth in LB was last recorded at 37°C and 15°C respectively, but their proportions of metabolically-active cells in the culture dropped below the 1% threshold at the immediately higher temperature points tested, i.e. at 42°C and 28°C respectively (Figs. 2 and 4).

**Fig. 6.**
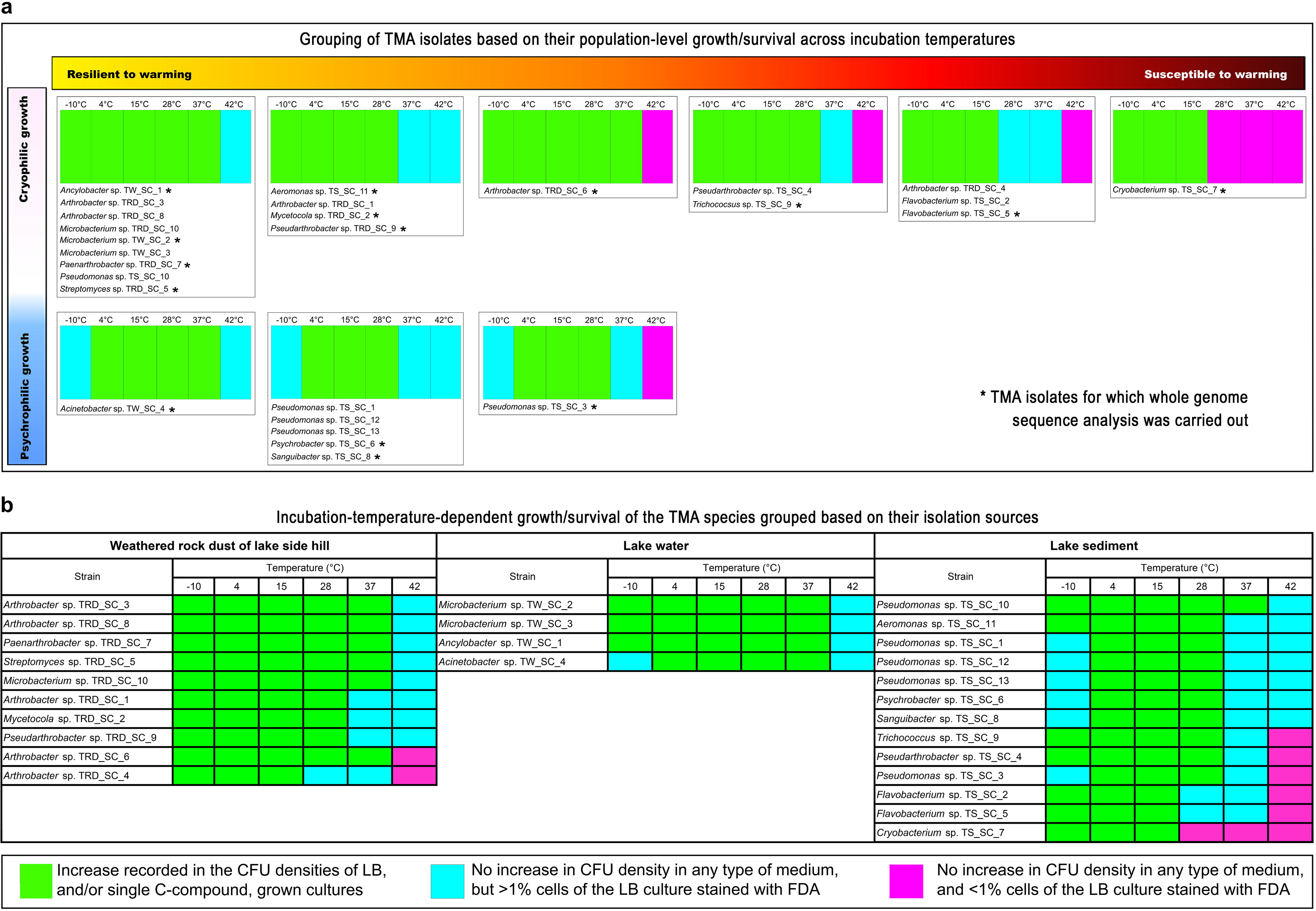
Graphic representation of growth, survival, or abolition of cell population for different TMA isolates after incubation in Luria broth for 28, 14, 2, 4, 2, and 1 days, at −10°C, 4°C, 15°C, 28°C, 37°C, and 42°C, respectively. (**a**) TMA isolates clustered based on the patterns of their responses to different incubation temperatures in terms of growth, survival, or abolition of cell population. (**b**) TMA isolates clustered on the basis of their isolation source.

So far as the most suitable temperature for LB-dependent growth was concerned (Fig. 5), 22 TMA isolates had their highest rates of increase in CFU density at 15°C. For all these isolates, except *Microbacterium* TRD_SC_10, 15°C growth rates exceeded their 4°C growth rates by orders of 10. Furthermore, for one half of the 22 TMA isolates having growth maxima at 15°C - i.e. for three *Arthrobacter* species, the two *Flavobacterium* species, and one species each of *Cryobacterium*, *Microbacterium*, *Pseudarthrobacter*, *Pseudomonas*, *Sanguibacter* and *Trichococcus* - 4°C growth rates were higher than their 28°C growth rates. In contrast, for the other half of these 22 isolates - i.e. for two *Arthrobacter* species, four *Pseudomonas* species, and one species each of *Aeromonas*, *Microbacterium*, *Paenarthrobacter*, *Pseudarthrobacter* and *Psychrobacter* - 28°C growth rates were higher than their 4°C growth rates. For *Acinetobacter* TW_SC_4, *Ancylobacter* TW_SC_1, *Microbacterium* TW_SC_2 and *Streptomyces* TRD_SC_5, maximum growth rates were recorded at 28°C. While the 15°C growth rates of all these four species were higher than their 4°C growth rates, for *Microbacterium* TW_SC_2 and *Streptomyces* TRD_SC_5 their 4°C growth rates were higher than the 37°C growth rates, but for *Acinetobacter* TW_SC_4 and *Ancylobacter* TW_SC_1 their 37°C growth rates were higher than the 4°C growth rates. For *Mycetocola* TRD_SC_2 alone, growth rate was highest at 4°C, then decreased through 28°C, and became negative at 37°C.

### Antibiosis by TMA actinobacteria

Of the 15 actinobacterial species isolated from the Tso Moriri area, 11 had the ability to inhibit the growth of at least one of the six higher-temperature-adapted foreign bacteria against which antibiosis was tested, namely the Gram negative organisms *Escherichia coli* K-12, *Advenella kashmirensis* WT001, *Paracoccus* sp. SMMA_5, and the Gram positive *Bacillus subtilis* SC_1, *Bacillus licheniformis* PAMA2_SD1, and *Lysinibacillus fusiformis* LAPE1_SD1 (Fig. 7). Only the four *Arthrobacter* species represented by the strains TRD_SC_1, TRD_SC_3, TRD_SC_4 and TRD_SC_8 had no antagonistic activity against any of the six target organisms. In contrast, *Pseudarthrobacter* TRD_SC_9 and *Streptomyces* TRD_SC_5 disallowed the growth of all the six foreign bacteria targeted, while *Paenarthrobacter* TRD_SC_7 inhibited all but *Advenella kashmirensis* WT001, and *Arthrobacter* TRD_SC_6 deterred only *Lysinibacillus fusiformis* LAPE1_SD1. On the flip side of the above data, most number of TMA actinobacteria were active against *Lysinibacillus fusiformis* LAPE1_SD1, *Bacillus licheniformis* PAMA2_SD1, and *Escherichia coli* K-12, which in turn were inhibited by 11, nine and seven actinobacteria respectively. Five, four, and three TMA actinobacteria were found to inhibit the growth of *Bacillus subtilis* SC_1, *Advenella kashmirensis* WT001, and *Paracoccus* SMMA_5, respectively. Notably, none of the 12 non-actinobacterial species isolated from TMA had the ability to inhibit the growth of any of the foreign bacteria against which antibiosis was tested (Fig. 7).

**Fig. 7.**
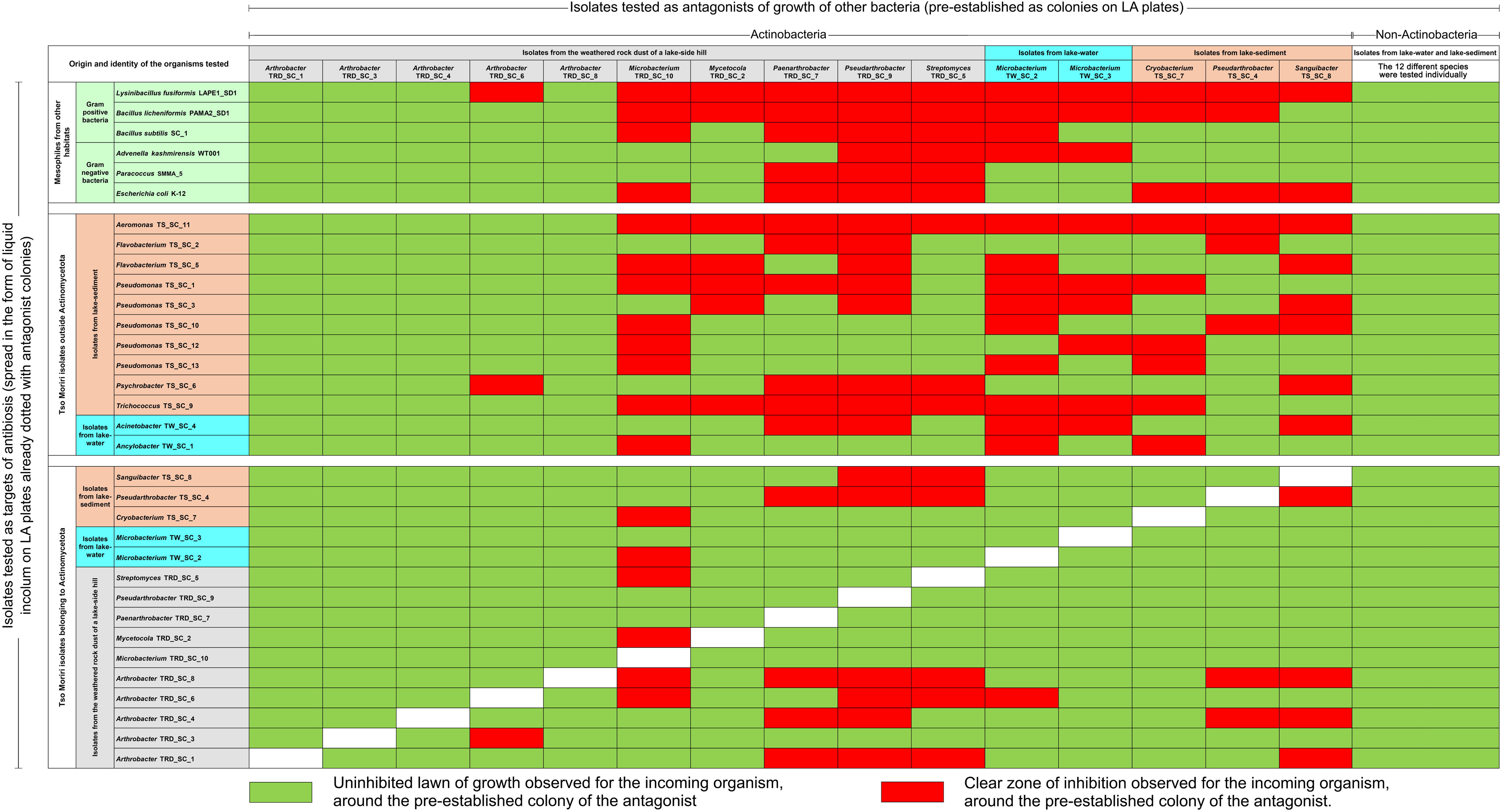
Graphic representation of the antibiosis potentials of the 27 bacterial species isolated from the Tso Moriri area. Green cells indicate growth, while red cells indicate inhibition of growth, for the target organisms.

The same 11 actinobacterial isolates, which had inhibited the growth of foreign mesophilic bacteria, also inhibited at least one or more non-actinobacterial TMA isolate(s) (Fig. 7). *Microbacterium* TW_SC_2 could inhibit the most number of TMA bacteria (nine) outside Actinomycetota. In contrast, *Arthrobacter* TRD_SC_6 could inhibit only one TMA bacterium outside Actinomycetota. From the reverse perspective, every non-actinobacterial TMA isolate was inhibited by at least three TMA actinobacteria. *Aeromonas* TS_SC_11 was the most vulnerable TMA bacteria outside Actinomycetota as its growth was inhibited by 11 TMA actinobacteria. In contrast, *Ancylobacter* TW_SC_1, *Flavobacterium* TS_SC_2, *Pseudomonas* TS_SC_12 and *Pseudomonas* TS_SC_13 were least prone to antibiosis as each of them was susceptible to only three actinobacterial isolates.

Out of the 15 TMA species belonging to Actinomycetota, eight could inhibit the growth of fellow actinobacterial isolates (Fig. 7). Overall, *Microbacterium* TRD_SC_10 and *Pseudarthrobacter* TRD_SC_9 inhibited the most number of actinobacteria (six each). In contrast, *Arthrobacter* TRD_SC_6, and *Microbacterium* TW_SC_2, inhibited only one TMA actinobacteria each. From the opposite point of view, 11 TMA actinobacteria were prone to inhibition by isolates belonging to the same phylum (Fig. 7). *Arthrobacter* TRD_SC_8 (inhibited by six TMA actinobacteria) was the most vulnerable TMA isolate belonging to Actinomycetota. In contrast, *Microbacterium* TW_SC_3, *Pseudarthrobacter* TRD_SC_9, *Paenarthrobacter* TRD_SC_7 and *Microbacterium* TRD_SC_10, were not prone to inhibition by any of the actinobacterial isolates.

In the context of antibiosis it was further noteworthy that none of the non-actinobacterial species isolated from the TMA could inhibit the growth of any other TMA isolate, whatever may be its phylum affiliation (Fig. 7).

### Key attributes of the whole genomes sequenced for selected TMA isolates

Complete whole genome sequence was determined for selected isolates from across the three environments explored within the Tso Moriri lake-desert ecosystem (Table 1). While one genome was sequenced from each of the 15 genera across which the 27 species-level isolates were classified, it was also ensured that at least one strain was analyzed from each cluster that had formed on the basis of growth or population-level survival at different incubation temperatures (Fig. 6a).

*De novo* hybrid assembly of the short and long DNA sequence reads generated using two different technologies yielded complete or near-complete genome sequences for all the TMA isolated analyzed (Tables 1 and S37). For 10 out of the 15 TMA isolates selected, their complete genomes were encompassed in single circular chromosomes: *Arthrobacter* TRD_SC_6, *Cryobacterium* TS_SC_7, *Flavobacterium* TS_SC_5, *Microbacterium* TW_SC_2, *Mycetocola* TRD_SC_2, *Paenarthrobacter* TRD_SC_7, *Pseudarthrobacter* TRD_SC_9, *Pseudomonas* TS_SC_3, *Sanguibacter* TS_SC_8 and *Trichococcus* TS_SC_9. For two TMA isolates their complete genomes were incorporated in single circular chromosomes plus multiple circular plasmids: *Acinetobacter* TW_SC_4 had two such plasmids of 18.1 kb and 81.9 kb length, while *Psychrobacter* TS_SC_6 had three of them, 6.4 kb, 7.1 kb and 23.1 kb in length. For *Aeromonas* TS_SC_11, *Ancylobacter* TW_SC_1 and *Streptomyces* TRD_SC_5, their near-complete genomes encompassed one or two uncircularized chromosomes plus one or two circular plasmids varying between 43.2 kb and 89 kb in length.

### Gene contents of TMA isolates concerned with adaptation to temperature extremes

Inspection of the eggNOG-annotated protein-coding gene sequence (CDS) catalogs of the 15 TMA isolates (Tables S38-S52) revealed diverse genes concerned with low and high temperature adaptations (Figs. 8 and 9a; Table S53). Besides a number of cold-adaptation-related genes concerned with cold-shock response and RNA remodeling, membrane fluidity regulation, and genome maintenance at low temperatures, several such genes were also detected which conferred tandem adaptation to low as well as high temperatures: these encoded chaperones and other protein quality control components, elicited DNA and oxidative stress responses, coded for global transcriptional regulators, or governed biosynthesis and transport of compatible solutes and osmoprotectants.

**Fig. 8.**
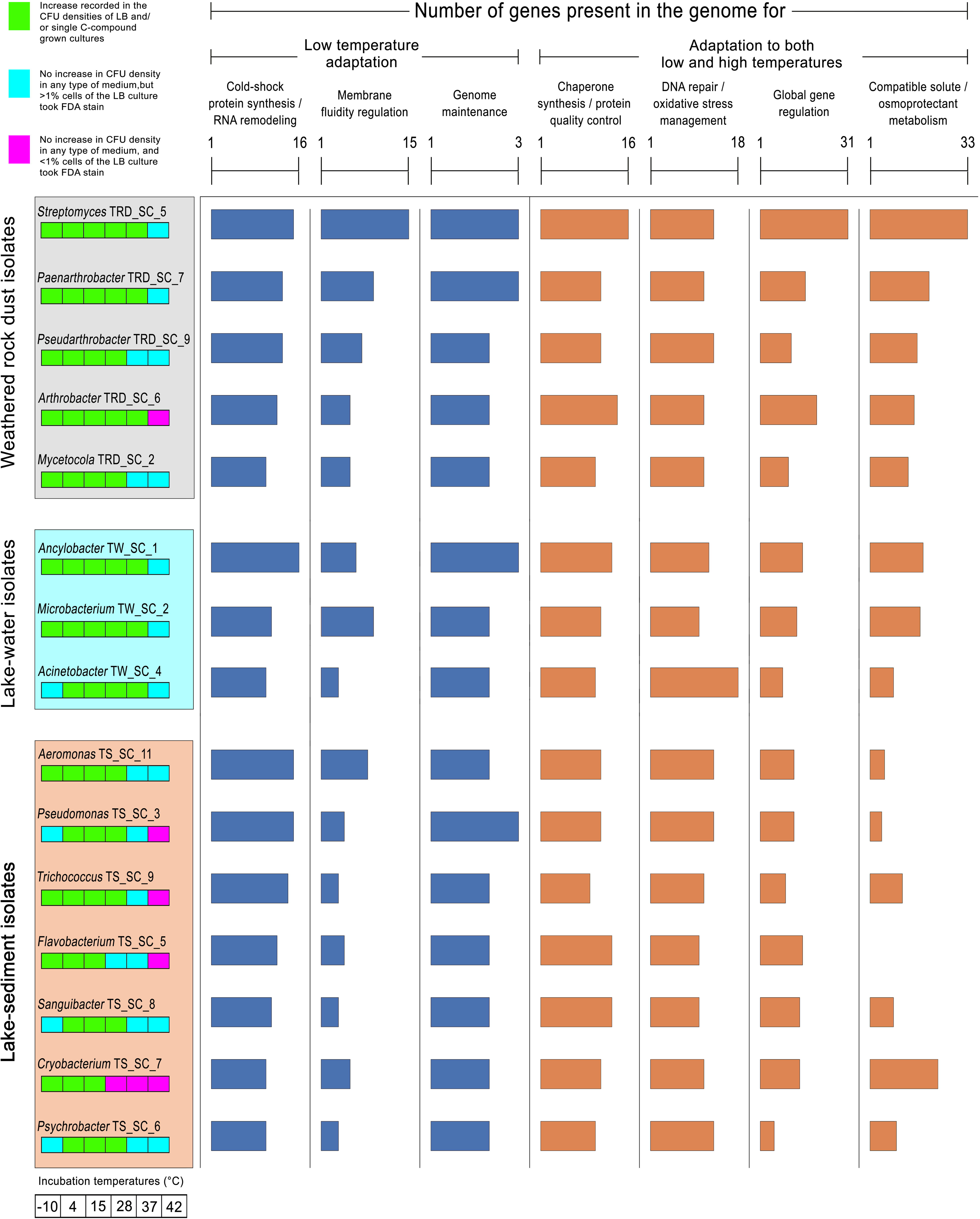
Metabolic-category-wise numeral distribution of genes putatively concerned with adaptations to low and high temperatures, within the genomes of the selected TMA isolates. Deep blue bars of the histograms indicate the numbers of genes associated with low temperature adaptation; orange bars indicate the numbers of genes associated with adaptation to low as well as high temperatures. Strains isolated from weathered rock dust, lake-water, and lake-sediment are clustered in boxes shaded grey, cyan, and light orange, respectively. Below each isolate name, a box panel indicating the organism’s growth/survival phenotype at the six incubation temperatures tested has been given (the fill color code of the boxes is given at the upper left corner of the figure).

**Fig. 9.**
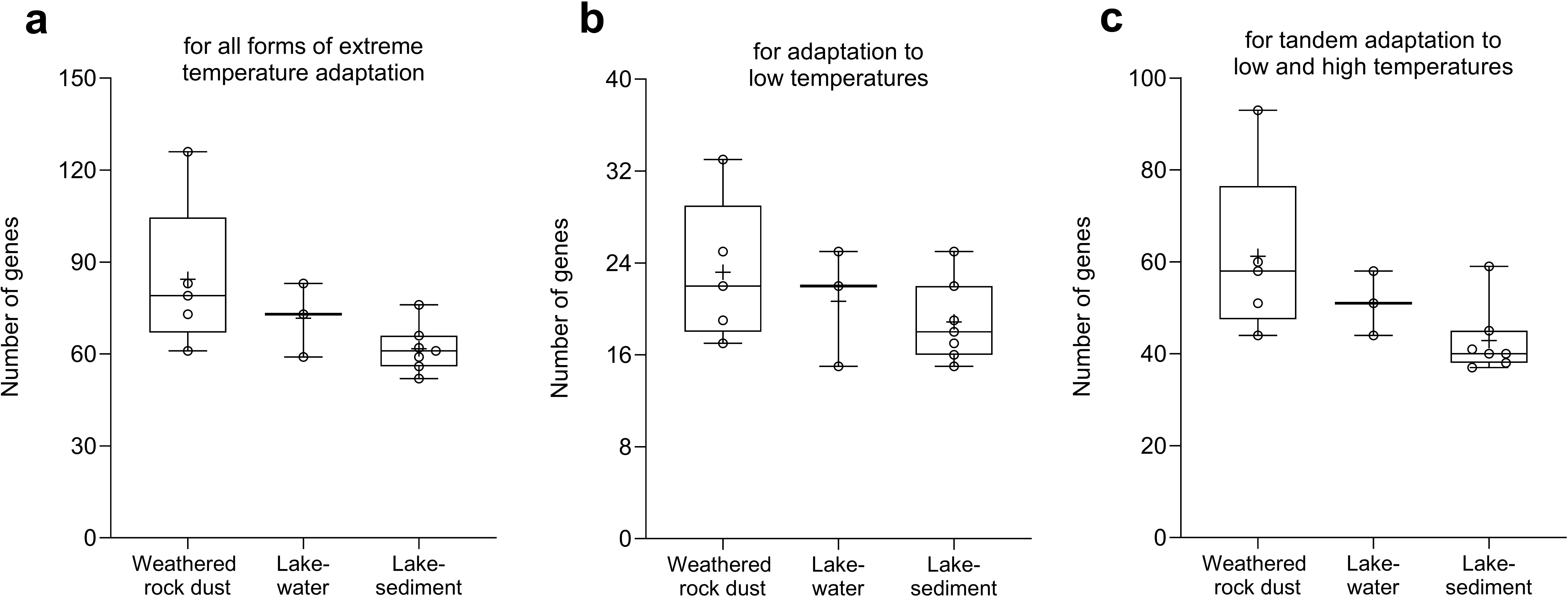
TMA-isolate-wise numeral distribution of genes putatively concerned with (**a**) all forms of extreme temperature adaptation, (**b**) low temperature adaptation, and (**c**) tandem adaptation to low and high temperatures (central tendencies of the data, post clustering of the strains according to their isolation sources, have been shown). Each isolation-source-specific boxplot displays the mean value (indicated by “+” sign), median value, interquartile range, and minimum–maximum whiskers, determined for the gene counts (indicated by hollow circles) of the isolates concerned.

*Streptomyces* TRD_SC_5 had the highest number of cold-adaptation-related genes [33], whereas *Acinetobacter* TW_SC_4 and *Psychrobacter* TS_SC_6 had the lowest [15]. So far as the genes involved in dual adaptation to low as well as high temperatures were concerned, *Streptomyces* TRD_SC_5 [93] and *Psychrobacter* TS_SC_6 [37] had the highest and lowest numbers of them, respectively.

The species isolated from weathered rock dust contained slightly higher diversities of genes [17-33 in number] for cold adaptation (Fig. 9b), compared with the isolates from lake-water [15-25 genes] and lake-sediment [15-25 genes]. Specifically, the *desKR* (KEGG orthology numbers: K07778 and K07693) two-component regulatory system, which controls membrane fluidity amid decreasing temperature (via modulation of fatty acid desaturase gene expression), was possessed mostly by rock-dust isolates (Table S53).

The species isolated from weathered rock dust also possessed relatively higher diversity of genes [44-93 in number] for dual adaptation to low and high temperatures, compared with the lake-water and lake-sediment isolates which possessed 44-58 and 37-59 such genes respectively (Fig. 9c). Specifically, the *treYZ* genes (K06044 and K01236) synthesizing the heat or cold, desiccation, and oxidative stress protectant trehalose; the *opuA*, *opuBD* and *opuC* genes (K05847, K05846 and K05845) governing the uptake of compatible solutes from the environment; and the *ectABC* genes (K06718, K00836 and K06720) synthesizing the protective compatible solute ectoine were possessed almost exclusively by the rock-dust isolates (Table S53).

### Genes encoding carbohydrate-active enzymes (CAZymes)

When the CDS catalogs of the selected TMA isolates (Tables S38-S52) were searched against the dbCAN3 and CAZy databases, a wide array of carbohydrate-active enzymes concerned with polysaccharide binding, modification, and degradation were revealed (Table S54). Amid a heterogeneous and overlapping distribution of genes across species isolated from the three TMA habitats, most number of putative CAZymes [293] was encoded by the genome of *Streptomyces* TRD_SC_5, whereas the least number of them [81] was encoded by *Psychrobacter* TS_SC_6. Overall, the detected CAZymes belonged to six broad functional categories - glycoside hydrolase (GH), glycosyl transferase (GT), carbohydrate esterase, polysaccharide lyase, enzymes with auxiliary activities, and carbohydrate-binding module (Fig. 10a). Of the different classes, again, GH and GT accounted for the most number of genes in all the genomes analyzed (collectively, 67-86% of all CAZyme genes possessed by the different strains were GHs and GTs). That said, most of the TMA isolates that were rich in GHs were relatively poorer in GT diversity, and vice versa. Only *Streptomyces* TRD_SC_5 had equal number of GH and GT genes, while *Microbacterium* TW_SC_2, *Arthrobacter* TRD_SC_6 and *Mycetocola* TRD_SC_2 also had comparable numbers of GH and GT genes. Overall, no clear environment- or habitat-specific trend was apparent when the species-wise distribution of CAZyme genes was considered in the context of the isolation source of the species (Fig. 10b).

**Fig. 10.**
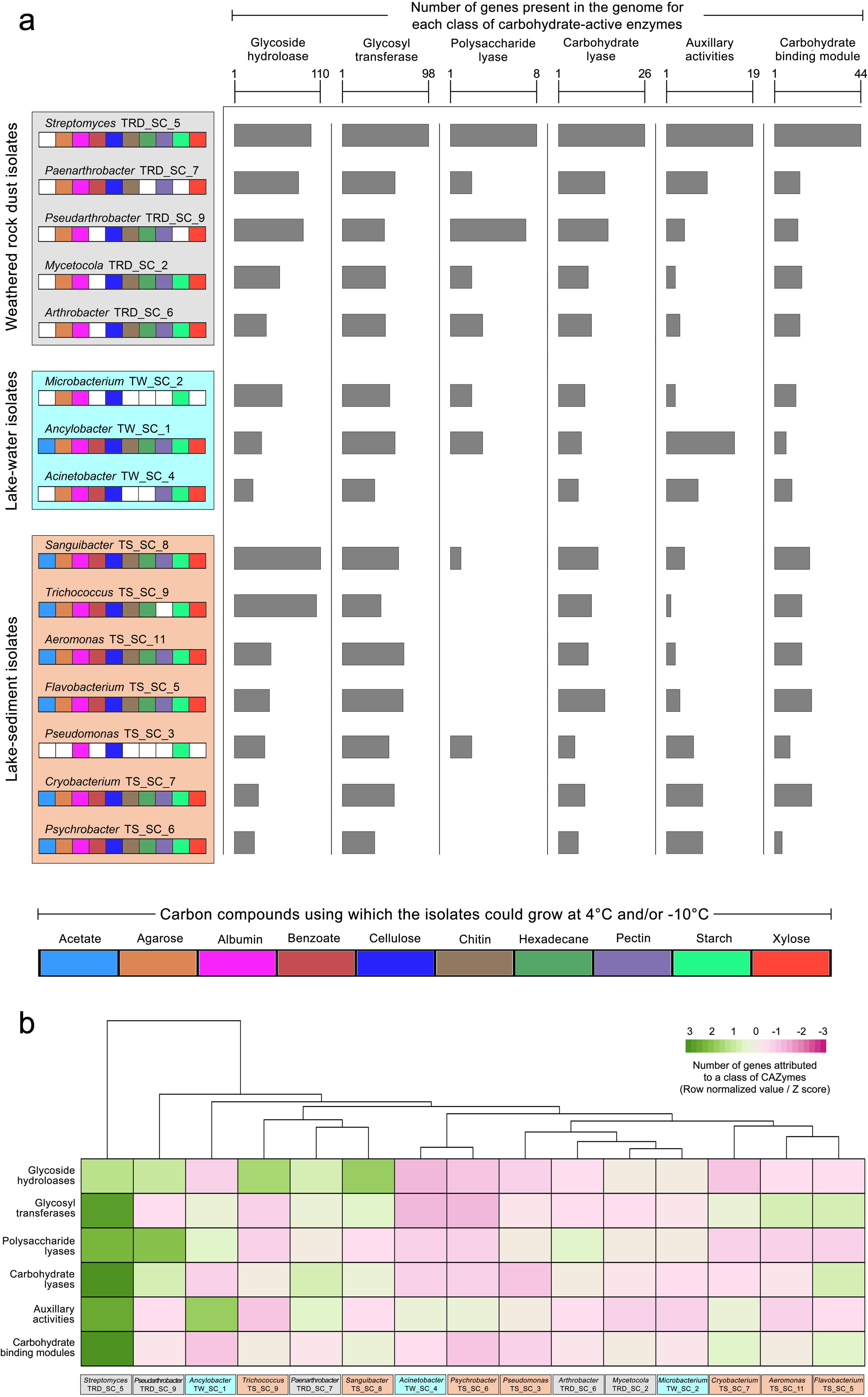
Functional-category-wise numeral distribution of genes putatively concerned with carbohydrate metabolism, within the genomes of the selected TMA isolates. (**a**) Representation of the data in the form of histograms, where grey colored bars indicate the numbers of genes encoding the different categories of carbohydrate-active enzymes (CAZymes). Strains isolated from weathered rock dust, lake-water, and lake-sediment are clustered in boxes shaded grey, cyan, and light orange, respectively. Below each isolate name, a box panel indicating the organism’s ability to grow using different carbon compounds (at 4°C and/or −10°C) has been given (the fill color code of the boxes is given below the figure). **(b)** Heatmap representing the functional-category-wise distribution of CAZyme-coding genes in the different TMA isolates (one-way hierarchical clustering of the isolates has also been rendered).

### Genes encoding antibiotics and other secondary metabolites

Overall, five classes of antibiotics [(i) penicillins and cephalosporins, (ii) prodigiosins, (iii) staurosporines, (iv) tetracyclines and (v) vancomycins; Table S55], and nine classes of secondary metabolites [(i) NI-siderophores, i.e. siderophores independent of non-ribosomal peptide synthase (NRPS), (ii) NAPAAs, i.e. non-alpha poly-amino acids, (iii) Class-II lanthipeptides, (iv) Class-III lanthipeptides, (v) NRPs, or non-ribosomal peptides, (vi) NRP-metallophores, (vii) terpenes, (vii) Type-I polyketides, and (ix) Type-III polyketides, all of which can potentially act as antimicrobial agents (Table S56)] were putatively synthesized by the 15 TMA isolates for which whole genome sequences were analyzed (Table 2).

**Table 2.**
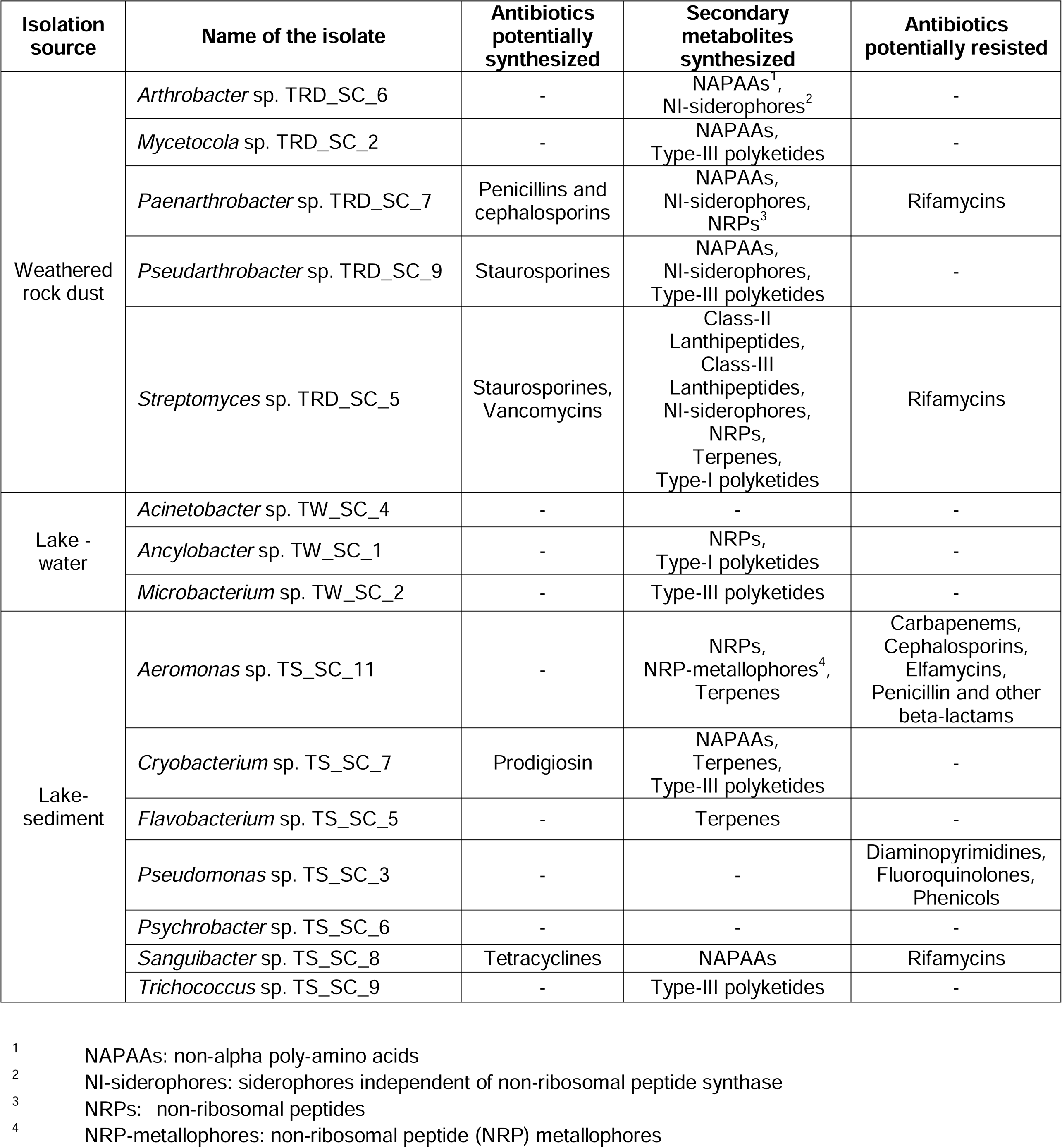
Genomic potentials of the TMA isolates for the synthesis of, and resistance against, different classes of antibiotic and secondary metabolite.

Every actinobacterium that was capable of inhibiting the growth of one or more target organism(s), and for which the complete whole genome sequence was analyzed, apparently possessed the ability to synthesize one or more classes of known antibiotic(s) and/or secondary metabolite(s). Among all the 15 isolates for which genomes were analyzed, highest diversity of antibiotics and secondarily metabolites was putatively synthesized by the actinobacterium *Streptomyces* TRD_SC_5 (this organism possessed key genes for the synthesis of Class-II and Class-III lanthipeptides, non-ribosomal peptides, siderophores independent of non-ribosomal peptide synthase, staurosporines, terpenes, Type-I polyketides and vancomycins). *Cryobacterium* TS_SC_7 (prodigiosins, NAPAAs, terpenes, and Type-III polyketides), *Paenarthrobacter* TRD_SC_7 (NAPAAs, NI-siderophores, NRPs, and penicillins and cephalosporins), and *Pseudarthrobacter* TRD_SC_9 (NAPAAs, NI-siderophores, staurosporines, and Type-III polyketides) possessed key genes for the synthesis of the next highest diversities of antibiotics and secondarily metabolites. Incidentally, these four actinobacteria had inhibited the growth of 14, 9, 15 and 20 out of the total 33 allochthonous and autochthonous microorganisms against which antibiosis was tested (Fig. 7). *Microbacterium* TW_SC_2 did not have genes required for the synthesis of any known antibiotic or secondary metabolite other than Type-III polyketides, but it had inhibited the growth of 14 target organisms (Fig. 7), presumably by hitherto unknown antimicrobial agents. On the flip side of the above data, no known antibiotic or secondarily metabolite was apparently synthesized by *Acinetobacter* sp. TW_SC_4, *Pseudomonas* sp. TS_SC_3 and *Psychrobacter* sp. TS_SC_6, which also had shown no antibiosis against any of the 33 targets tested. *Aeromonas* TS_SC_11, *Ancylobacter* TW_SC_1, *Flavobacterium* TS_SC_5, and *Trichococcus* TS_SC_9 too had not inhibited any target organism; corroboratively, they did not also synthesize any known antibiotic (the four species, however, putatively synthesizied a few classes of secondary metabolites; Table 2).

### Genes conferring potential resistance against antibiotics

Of the 15 isolates for which genomes were analyzed, only five were found to posssess genes central to resistance against carbapenems, cephalosporins, diaminopyrimidines, elfamycins, fluoroquinolones, penicillins and other beta-lactams, phenicols, rifamycins, teicoplanins, and/or vancomycins (Tables 2 and S57). The remaining 10 genomes were found to encompass no such gene which is central to resistance against any known antibiotic. While potential resistance against rifamycins was accounted for by the highest number of genomes (three), putative resistance against highest number of antibiotic groups was possessed by *Aeromonas* TS_SC_11. This isolate, however, was inhibited by several actinobacterial isolates, apparently by antimicrobials other than penicillins or other beta-lactams, carbapenems, cephalosporins and elfamycins, against which TS_SC_11 possessed putative resistance according to the genomic data (Fig. 7; Table 2).

## DISCUSSION

### Habitat as the key driver of adaptation to extremes of temperature

All the genera under which the TMA isolates were classified, excepting *Ancylobacter* and *Paenarthrobacter*, encompassed multiple members retrieved previously from other natural or artificial cold/frigid environments (Table S58; also see Supplementary References). Remarkably, each of them also encompassed at least one member retrieved previously from a mesic/hot habitat (Table S59; also see Supplementary References). Consistent with these facts, cell populations of different TMA isolates belonging to the same genus were found to exhibit differential growth/survival at a given incubation temperature (Fig. 6a).

Native environments of the TMA isolates, rather than their phylogenetic background, seemed to exert the most crucial influence on adaptation to extremes of temperature. For all the genera isolated in this study, except *Paenarthrobacter*, member strains from other parts of the world, especially mesic/hot environments, are known to grow at temperatures higher than the maxima recorded for the TMA counterparts (Fig. S5; Table S60; also see Supplementary References). As for *Paenarthrobacter*, it was intriguing as to why the growth temperature maxima of its strains reported earlier from warmer habitats, and that of the current isolate from cold/frigid environment, converged at 37°C. It was, however, understandable that the growth temperature minimum of *Paenarthrobacter* (Fig. S5; Table S60) was pegged thus far at 4°C (42, 43) because no member of this genus had been isolated from any cold habitat before this study. With regard to variable cold adaptation in closely-related bacteria, it was further noteworthy that none of the genera isolated from Tso Moriri area, except *Cryobacterium*, *Pseudomonas*, *Psychrobacter* and *Sanguibacter* (44–47), encompassed any such strain from any other part of the world which possessed growth temperature minimum lower than its TMA counterpart (Fig. S5; Table S60).

Certain common phenotypes seemed to unify majority of strains isolated from a given TMA environment (Fig. 6b).

1. All the bacteria isolated from the weathered rock dust of the lake-adjoining hill, with the exception of only *Arthrobacter* TRD_SC_4 and TRD_SC_6, grew or survived at the population-level through the entire temperature range tested (i.e. −10°C to 42°C).
2. The bacteria from Tso Moriri’s water also accomplished population-level growth or survival through the entire range of incubation temperatures tested.
3. Most strains isolated from Tso Moriri’s sediment had population-level growth/survival across narrower ranges of incubation temperature, compared to the isolates from rock-dust and lake-water.

Concurrent with the above phenotypic trends, cumulative diversity of genes concerned with all forms of extreme temperature adaptation was relatively higher in the rock-dust and lake-water isolates, compared to those from the lake-sediment (Fig. 9a; Table S53). Individually also, the diversity of genes concerned with adaptation to low (Fig. 9b; Table S53), and low as well as high (Fig. 9c; Table S53), temperatures were relatively higher in the rock-dust and lake-water isolates, compared to those from the lake-sediment.

Tso Moriri’s surficial water freezes in the winter and thaws in the summer. Likewise, the rocky slopes of the adjoining hills get covered by snow in the winter and the same melts through the summer; furthermore, the rocks are exposed to extreme heating and cooling on a diurnal basis. Thus, wider temperature windows for population-level growth/survival, and enrichment of corresponding genetic resources, in bacteria from these two environments could be reflective of a positive linkage between adaptation to the extremes of temperature and the variability of the thermal condition prevailing in the habitat. By the flip side of the same reasoning, it was also quite natural for the bacteria from Tso Moriri’s subsurface to have narrower temperature windows (and genes for all forms of extreme temperature adaptation) for population-level growth/survival, compared to their surficial neighbors.

In the context of environment as a key driver of low temperature adaptation, it was further noteworthy that for *Acinetobacter*, *Aeromonas*, *Microbacterium*, *Paernarthrobcter*, *Pseudarthrobacter* and *Trichococcus*, strains known to grow at genus-specific minimum temperatures before this study (Fig. S5; Table S60) were all isolated from mesic or hot environments (42, 48–52), even though other strains of all these genera except *Paenarthrobacter* had been isolated from natural and/or artificial cold/frigid environments (Table S58). On the flip side, strains of *Aeromonas* and *Mycetocola* that were known to grow at genus-specific temperature maxima before this study (Fig. S5; Table S60) were all isolated from natural or artificial cold/frigid environments (53, 54), even though other strains of these genera had been isolated from mesic or hot environments (Table S59). These habitat-phenotype mismatches could be the upshots of drastic microbial transportation across physicochemically distinct ecosystems, alongside the existence of some hitherto unappreciated metabolic convergence between high and low temperature adaptations. Such a counterintuitive idea was reinforced by the facts that out of the 20 cryo-adapted (−10°C-growing) bacteria isolated from the TMA, 16 and 10 could grow at 28°C and 37°C respectively (Fig. 6a). Furthermore, after 1 day at 42°C, only eight TMA isolates had >0.1% CFU left in the culture (Figs. 4f and 4i), and among these, six could also grow at −10°C (Fig. 2b).

### Autochthonous populations of heat-susceptible psychrophiles as biogeothermometers of *in situ* warming

TMA isolates, irrespective of their phylogenetic affinities, exhibited different levels of thermal susceptibility. For instance, almost all cells of *Cryobacterium* TS_SC_7 lost their divisibility, as well as metabolic activity, after a four-day exposure to 28°C (Figs. 4b and 4g), so their retrieval from the lake’s sediment could be indicative of the fact that the temperature of this under-water substratum either does not reach 28°C, or even if it does so, it does not remain at that level for several days. On the flip side, elimination of such bacteria from environments where they had been prevalent previously would indicate protracted *in situ* warming up to and above 28°C. Population dynamics of TS_SC_7-like bacteria can, therefore, be construed and contrived as spatiotemporal biogeothermometers of cold/frigid environments that are indisposed to prolonged physical monitoring.

Almost all cells of *Flavobacterium* TS_SC_5 and TS_SC_2 lost their metabolic activity after a full day at 42°C *in vitro* (Figs. 4f and 4i), while their divisibility was already abolished after a single day’s exposure to 37°C (Fig. 4c), or even after a two-day exposure to 28°C (Fig. 4a). Likewise, almost all cells of *Arthrobacter* TRD_SC_4 lost their activity after a full day at 42°C (Figs. 4f and 4i), while their divisibility was already abolished after a two-day exposure to 37°C (Figs. 4d and 4h), and only a quarter of the cell population retained its divisibility after four days at 28°C (Figs. 4b and 4g). Thus, a progressive decline in the prevalence of a natural population of *Flavobacterium* species similar to TS_SC_5 and TS_SC_2, over time and/or space, can be correlated with a continuing temporal and/or spatial escalation of temperature to 28°C and above. The same for TRD_SC_4-like populations can, in turn, corroborate ongoing spatiotemporal increases of *in situ* temperature to 37°C and above. In the same vein, population dynamics of bacteria like *Arthrobacter* TRD_SC_6 and *Trichococcus* TS_SC_9, within cold/frigid ecosystems, can be used as real-time bellwethers signaling shifts of *in situ* temperature towards 42°C.

### A baseline for *in situ* carbon remineralization under cold/frigid conditions

High levels of organotrophic growth were recorded for all the TMA isolates at 4°C; slow but measurable growth also took place for ∼75% of them at −10°C (Figs. 2 and 3). How these *in vitro* metabolic abilities translate into community-level actions *in situ* is central to our knowledge on the baseline of organic matter degradation (carbon remineralization) within the cryosphere. Although biogeochemical data are not yet available for the Tso Morir ecosystem, permeating presence of bacteria similar to the species analyzed genomically was revealed across the three TMA habitats. Small but definite proportions of metagenomic reads obtained from Tso Moriri’s water and sediment samples, as well as the sample of weathered rock dust from the lake-side hill, mapped onto each of the 15 whole genomes used as target for sequence alignment. Collectively, the 15 genomes accounted for 0.1%, 0.3% and 0.2% of metagenomic reads from the lake-water, lake-sediment and rock-dust samples respectively (Table S61). These numbers not only illustrated the ubiquity of cryo-adapted and cryo-tolerant copiotrophs similar to the few specimens analyzed here, but also envisaged a biogeochemically effective status for them, across the Tso Moriri ecosystem. Effectiveness of psychrophilic copiotrophs in the biogeochemical cycling (remineralization) of carbon across the Tso Moriri ecosystem was further corroborated by the wide array of carbohydrate-active enzymes detected in the genomes of the current isolates (Fig. 10; Table S54), as well as the assembled metagenomes of the lake-water, lake-sediment, and rock-dust samples (Tables S62).

### Thermal susceptibility of autochthonous organotrophs can usher negative feedback control/reversal of warming in cold/frigid ecosystems

Within cold/frigid ecosystems vulnerable to global warming and anthropogenic perturbations (55–57), microbial activity is thought to remain minimal as long as the temperature remains below or around 0°C. However, organic matter degradation, and consequent greenhouse gas emission, starts once *in situ* temperature rises and cryoturbation takes place due to season and/or climate change (58, 59). Increased greenhouse effect brought about by enhanced microbial activities stimulates the biotic processes all the more, and that in turn ushers further thawing via CO_2_ and CH_4_ emission. In this way, a positive feedback loop of cyclical warming (Fig. 11) is operationalized in the ecosystem (27, 32, 33).

**Fig. 11.**
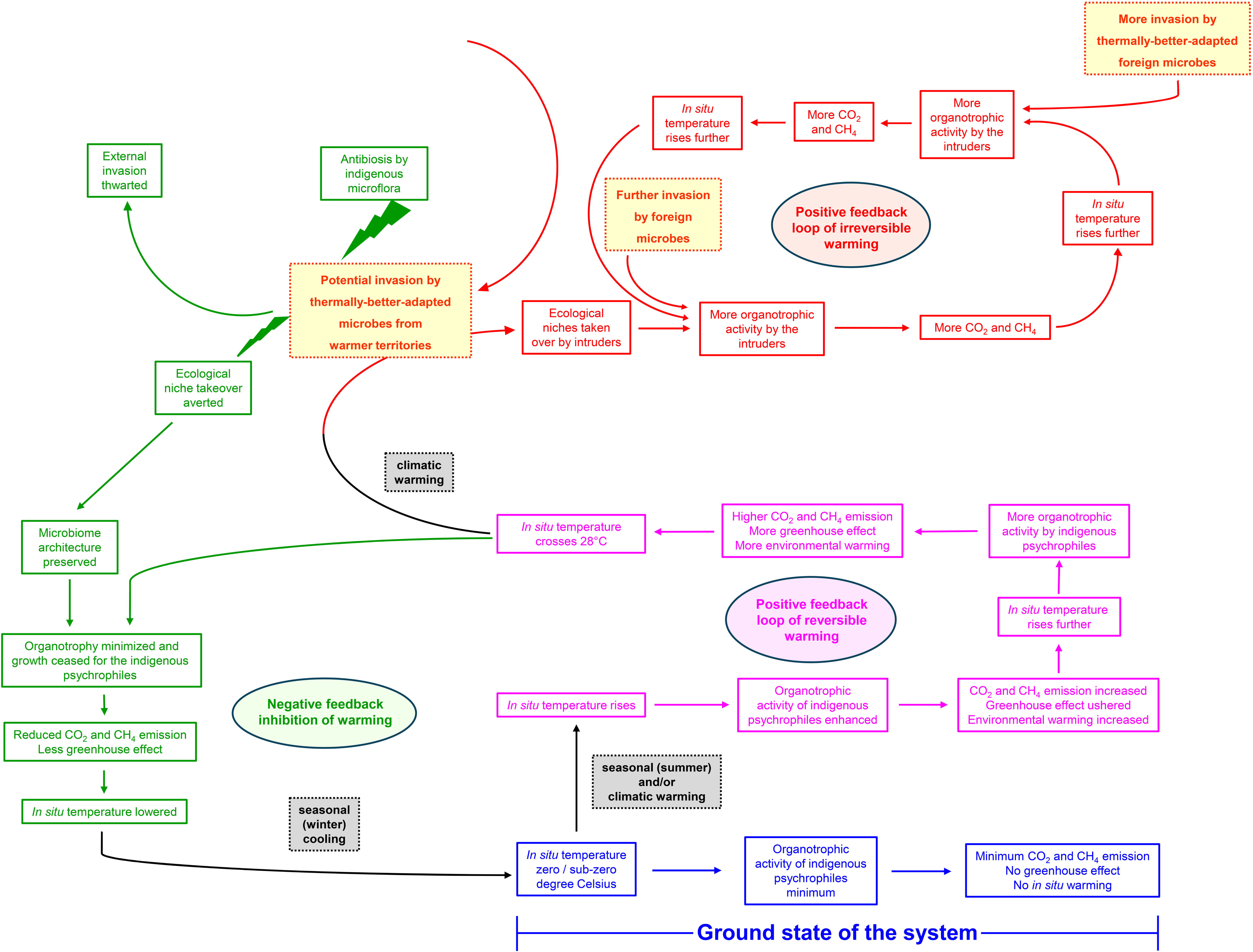
Schematic diagram showing how indigenous psychrophilic bacteria might render homeostatic control of *in situ* temperature in the context of climate warming and consequential threat of invasion by foreign microbes from warmer territories.

Psychrophilic (4-15°C) growth proficiencies recorded for all the TMA isolates *in vitro*, together with the growth potentials exhibited by most of them at 28°C, indicated that, in nature, microbes-mediated positive feedback cycles can abet environmental warming (via greenhouse gas generation) within the 4°C to 28°C range. On the other hand, above 28°C, gradual cessation of organotrophic growth and activity (Fig. 4) can cut back on the *in situ* production of simple fatty acids and CO_2_, which constitute the central substrates of methanogenesis, the terminal process of the carbon cycle (60). Termination of organotrophy, followed by the preemption of methanogenesis, curbs the emission of greenhouse gases, triggers a negative feedback control of warming, and usher over time, a course reversal in the vicious cycle of “warming - microbial growth - and further warming”. Seasonal (winter-mediated) cooling eventually lowers the *in situ* temperature back to the zero and sub-zero degree Celsius levels (Fig. 11).

On the part of a cold/frigid ecosystem, exercise of negative feedback control (termination of autochthonous microbial growth and reduction of metabolic activity; see Fig. 11) towards the restoration of homeostatic balance (reversing the trends of *in situ* warming), is fraught with the danger of indigenous microbes being removed from the habitat, and their ecological niches taken over by more-thermotolerant intruders from discrete geographical territories (Fig. 11). Such incursions can rapidly transform microbiome architectures if not abated at temperatures below the critical level (61) where most cells of most species of the native microflora lose their growth as well as metabolic activity. Potential microbiome transformations can, in the long run, alter ecosystem structures and functions drastically, and in doing so tilt the temperature equilibrium irreversibly in favour of the positive feedback mechanism (27, 32, 33) which promotes warming (Fig. 11).

### Heat-enduring actinobacterial psychrophiles as defenders of cold/frigid microbiomes

In the scenario of an imminent, warming-mediated microbiome alteration, antibiosis potentials of heat-enduring actinobacteria native to the cold/frigid ecosystem (Fig. 7) can deter foreign microbes from colonizing the habitat and apprehending the ecological niches of the indigenous psychrophiles. In the long run, this kind of niche safeguard enhances the chances of population rejuvenation for all autochthonous cold-adapted species upon reversal of thawing (Fig. 11).

Actinomycetota members are known for their extraordinary abilities to produce secondary metabolites including antibiotics, which confer them adaptive fitness against diverse environmental challenges, resulting in the high ecological amplitude of the phylum (62–64). Moreover, they are often abundant in cryospheric microbiomes (65, 66), and are known to protect indigenous microbial communities from external invaders in other critical ecosystems (67). So far as the 15 TMA actinobacteria were concerned, a majority of them were substantially thermotolerant (Figs. 4 and 6a). On top of this, 11 TMA actinobacteria were capable of inhibiting *in vitro* Gram positive and/or Gram negative bacteria from discrete higher-temperature ecosystems at 28°C (Fig. 7), which is lower than the critical high temperature for most of the TMA isolates (Fig. 4). It is, therefore, not unlikely that in the face of plausible microbial incursions from warmer territories, native actinobacterial psychrophiles would thwart foreign occupation and protect indigenous microorganisms from being vanquished from the habitat. Furthermore, in this context it was noteworthy that among the 11 TMA actinobacteria exhibiting antibiosis potentials against foreign bacteria, six had been isolated from the rock-dust sample, while three and two were from the lake’s sediment and water respectively (Fig. 7). This showed that microbiome protection by antibiosis-enabled heat-enduring actinobacterial psychrophiles could be widespread across physicochemically distinct cold/frigid environments.

### Intricate niche partitioning as central to microbiome functioning within cold/frigid ecosystems

In relation to the broader ecological implications of antibiosis it was noteworthy that a number of TMA actinobacteria eliminated not only foreign microorganisms, but also co-inhabitants of their own environment as well as natives of their adjacent habitats (these included fellow actinobacteria also; see Fig. 7). These findings brought to the fore extensive niche partitioning (68, 69) as a plausible stratagem of microbiome functioning within the cold/frigid ecosystem.

However, neither the phenotypic data, nor the genome-based predictions, available at present in relation to antibiosis (and resistance against the same) could elucidate how ecological niche separation pans out in the micro-environment scale, within the Tso Moriri habitats. In other words, it was not possible from the present data to decipher how niche separation within the TMA habitats effectively precluded direct physical contacts between the actinobacterial antagonists and the species they inhibited *in vitro*. This limitation was attributed to the fact that bulk sampling had already smudged all imprints of niche differentiation, and deciphering of the same would require a new generation of very-finely-resolved spatial study of community ecology.

Genome-based predictions of the TMA isolates’ potentials for synthesizing and resisting different antibiotics and secondary metabolites were also not sufficient to elucidate the molecular mechanisms underlying the antibiosis phenotypes recorded *in vitro*. Nevertheless, genomic potentials of the TMA isolates for antibiotic and secondary metabolite synthesis (Table 2) were consistent with their phenotypes recorded in relation to pair-wise antibiosis (Fig. 7). Furthermore, the TMA isolates’ antibiotic resistance potentials predicted from genome contents (Table 2) contradicted neither the antibiosis phenotypes recorded *in vitro* nor the genotypes delineated with regard to antibiotics and secondary metabolites synthesis (Table 2). In a nutshell, for every instance where an isolate had inhibited the growth of another TMA species, the antagonist was found to have the genomic potential for synthesizing at least one such class of antibiotic or secondary metabolite against which the inhibited species had no resistance gene.

### Using TMA copiotrophs to address the concerns of biodegradation in the cryosphere

Besides global warming, burgeoning human activities have, in recent times, brought about unprecedented perturbations in the delicately balanced cold/frigid ecosystems that are dispersed across the vast high-altitude and high-latitude territories of the world. One of the major factors detrimental to these pristine ecosystems is the accretion of anthropogenic wastes. The pace of microbial growth (19, 26), and thereby degradation of organic matter (26, 70, 71), being slow at temperatures near and below the freezing point of water (also see Figs. 2 and 5), waste-treatment systems (bioreactors) operated in alpine and polar territories require external energy input for optimum-temperature maintenance (72); consequently, these contrivances are neither eco-friendly nor cost effective. Even bioreactors that employ microbes isolated from pristine cold/frigid habitats also sometimes fail to deliver efficient results because psychrophilic microorganisms, owing to their residence in, and adaptation to, organic-carbon-scarce habitats (31, 73), are often oligotrophic in nature (74, 75); accordingly, they are unfit for proliferation in the nutrient-rich environments of bioreactors. Moreover, tens of degree-Celsius thawing above the freezing point of water runs the risk of making psychrophilic bacterial populations uncompetitive against the thermotolerant, and mostly copiotrophic, microorganisms that find their way into the bioreactors along with the waste materials introduced for degradation (76, 77). In the face of all these challenges, microbes such as the TMA psychrophiles – by virtue of their abilities to (i) grow via catabolism of simple to complex carbon compounds over a wide range of temperature, (ii) grow under low as well as high concentrations of organic matter, and (iii) thwart the proliferation of thermotolerant microbes from warmer habitats – can become ideal constituents of bioreactors designed to perform through zero and sub-zero degree Celsius temperatures.

## MATERIALS AND METHODS

### Study site

The Himalayas and Trans-Himalayas encompass the largest reservoir of snow and ice on Earth outside the Arctic Circle and Antarctica (78). Within the cold arid vastness of the Himalayan rain shadow, the territory of Ladakh features an extensive high-altitude plateau, where multifaceted microbial life thrives to support homeostatically fragile ecosystems harboring unique multi-stress-adapted plants and animals (28, 79–82). Covering a large swathe of eastern Ladakh (Fig. 1a) sprawls the Changthang desert dotted by several small to large lacustrine bodies among which Tso Moriri is one of the most prominent (Fig. 1b). This gigantic water body (Figs. 1b-c), located at an altitude of 4522 m, is flanked on all sides by barren hills (Figs. 1d-e) that rise either from a distance or very close to the bank of the lake (Fig. 1d). Small patches of lush green meadows, marshes, and wetlands hem only the northern and southwestern shores of Tso Moriri where two big glacial streams pour their water into the lake. Rest of the cold-desert habitat around the lake is characterized by a bleak topography and highly inhospitable physicochemical conditions that are paralleled only by the extreme multi-stressor environments of a few high-altitude areas of the Chilean and Argentinean Andes (83, 84).

### Sampling

On 28 October 2021, water (Fig. 1f) and sediment (Fig. 1g) samples were collected from near the western shore of Tso Moriri (at GPS coordinates 32°94′ N and 78°27′ E), together with samples of weathered rock dust from the barren slope of a hill overlooking the western bank of the lake (Fig. 1e). The entire hill slope, rising from ∼100 m off the Tso Moriri shore, was devoid of any vegetative cover; visibly, not a single blade of grass grew on it.

In order to arrest the aquatic microbiota, as described previously (85), five different batches of 500 mL water were sampled from five discrete sites located at intervals of 1 m, and within 1 m from the shore of the lake. Each batch of 500 mL water was aspirated from within 1 cm of the lake surface using a sterile syringe (Tarsons Products Limited, India), and passed through an autoclaved, Swinnex holder-mounted (Merck KGaA, Germany), sterile, mixed cellulose ester membrane filter (Merck Life Science Private Limited, India) having a mesh size of 0.22 μm and diameter of 47 mm. Subsequent to filtration, each membrane corresponding to the cell residue of 500 mL lake-water was dislodged from the Swinnex holder, folded using autoclaved forceps, and inserted into a 7.5 mL autoclaved cryovial that contained 5 mL of 15% (v/v) glycerol supplemented with 0.9% (w/v) NaCl (86).

From the same five locations where Tso Moriri’s water was sampled, sediments were collected in approximately equal quantities, and pooled inside a single 250 mL polypropylene bottle. Inside the polypropylene bottle the sediment fractions were mixed thoroughly to give rise to a bulk sample which was used for all downstream investigations. At each sampling point, soft deposit was scraped carefully from the sediment-water interface using a flat and wide sterile spatula without disturbing layers deeper than 1 cm from the sediment-surface.

Dry and loose, weathered rock dust samples (fine talus and scree particles) were collected in approximately equal quantities from five distinct points situated within one meter from each other, on the lake-facing slope of the aforesaid hill. At each sampling site, fine powdery materials were scraped cautiously from the surface without disturbing layers deeper than the top 1 cm. The five sample fractions were pooled inside a 250 mL polypropylene bottle, and mixed thoroughly with a sterile spatula to yield a bulk sample that was used for all subsequent investigations.

After the insertion of a cell-precipitated membrane, or sediment / rock-dust sample, each cryovial or polypropylene bottle was capped tightly, sealed with parafilm (Tarsons Products Limited, India), and put inside a polyethylene bag, which eventually was packed in a heat-insulated ice box and shipped by air to the laboratory. All samples were investigated immediately upon their arrival at the laboratory.

In order to extract metagenomic DNA from Tso Moriri’s water and sediment, and also from the rock-dust of the lake-side mountain, sampling was carried out as described above. Only for the water sample a few minor adjustments were made. Total 5 L of the lake-water was passed through five separate (pre-autoclaved) 0.22 μm filters (1 L per filter), which in turn were inserted into five different sterilized cryovials, each containing 5 mL of sterile 50 mM Tris:EDTA (pH 7.8).

### Enrichment and isolation of cryo-tolerant / cryo-adapted, psychrophilic copiotrophs

In order to enrich copiotrophic, cryo-tolerant / cryo-adapted, psychrophiles from Tso Moriri’s water, each 0.22 μm cellulose acetate filter, which contained microbial cell residue from 500 mL lake-water, was shredded with sterile scissors inside the same vial in which it was inserted on-field. The vial was whirled for 15 minutes; after that the filter-shreds were allowed to settle at the bottom; finally, the supernatant was collected in a fresh sterilized vial without disturbing the debris. This process was repeated for all the five filter-containing cryovials involved in lake-water sampling, and the supernatants recovered from each of them were pooled to get an approximately 22 mL cell-suspension in NaCl-glycerol, which corresponded to 2.5 L of bulked lake-water. This ∼22 mL cell-suspension was added to 80 mL LB prepared in such a way that the actual nutrient concentrations were achieved after the mixing. The inoculum-medium mixture was subjected to three consecutive cycles of “7 day freezing at −10°C, followed by 7 day thawing at 4°C”, after which pure cultures were isolated at an incubation temperature of 4°C by means of dilution plating, picking of visibly-distinct single colonies, and repeated dilution streaking on Luria agar (LA) plates.

To enrich copiotrophic, cryo-tolerant / cryo-adapted, psychrophiles from the lake-sediment or rock-dust sample, 1 g of the corresponding material was added to 100 mL LB, and the suspension was subjected to three cycles of “7 day freezing at −10°C, followed by 7 day thawing at 4°C”. After the third round of thawing, pure culture strains were isolated at 4°C in the same way as described above.

All the new isolates were maintained in LA slants with a standard transfer interval of 15 days. For routine growth in LB, or fortnightly transfer in LA, all isolates were grown at 4°C. The strains were classified, as described previously (87), up to the lowest taxonomic rank that was ascribable based on 16S rRNA gene sequence similarities with validly-published species curated in the List of Prokaryotic names with Standing in Nomenclature (88).

### Determining the temperature window for growth

The temperature range over which the new isolates could, or could not, grow was delineated by recording the extent to which CFU density increased, or decreased, in the individual LB cultures of the strains, after aerobic incubation at −10°C, 4°C, 15°C, 28°C, 37°C, and 42°C. For a given experiment, a seed culture of the strain tested was prepared by transferring a loopful of cell mass from a 7 day old LA slant culture to fresh 20 mL LB medium kept in a 50 mL Erlenmeyer flask, which was then incubated aerobically at 15°C until when the culture attained its mid-log stage. From this 20 mL seed culture, 1% inoculum was transferred to fresh 20 mL LB medium to set up the test culture. Experiments checking aerobic growth at incubation temperatures ranging between 4°C and 42°C were carried out in 50 mL Erlenmeyer flasks, whereas those checking aerobic growth at −10°C were carried out in 50 mL polypropylene tubes.

To record CFU density at any time point, 1 mL of the experimental culture concerned was serially diluted using 0.9% (w/v) NaCl, and plated on LA in triplicates (in case of −10°C incubations, the frozen cultures were first liquefied via thawing at 4°C, and then subjected to dilution plating). Subsequently, the LA plates were incubated at 15°C, and colonies appearing on them were counted after 2-3 days depending on the growth rate of the culture in question. Finally, the CFU density was calculated by first multiplying the colony-counts of the individual dilution-plates by their corresponding dilution factors, and subsequently adding and averaging the values across the dilution grades and replica plates available.

The above experiments were repeated for the comparator organism *Escherichia coli* by recording the increases or decreases in CFU density that the LB cultures of strain K-12 underwent after incubation at −10°C, 4°C, 15°C, 28°C, 37°C, and 42°C. *E*. *coli* seed cultures were prepared in the same as those prepared for the TMA isolates except for the fact that incubations were carried out at 28°C for 12 h. Experimental cultures too were subjected to the same procedure as above, while CFU densities for K-12 were recorded by incubating the colony-counting LA plates at 28°C for two days.

When the final CFU density of an experimental culture recorded after the stipulated period of incubation was higher than its initial (0 hour) CFU density, the growth rate of the concerned isolate, at the temperature in question, was calculated as the percentage change that was recorded in the CFU density over time (percentage of the initial CFU mL^-1^ culture that increased day^-1^ incubation).

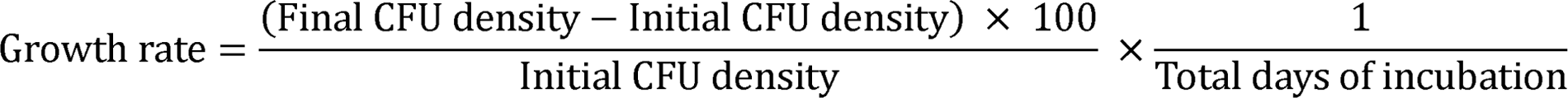

### Determining the temperature window for population-level survival

When the final CFU density of an experimental culture recorded after the stipulated period of incubation was lower than its initial (0 hour) CFU density, the survival frequency of the concerned isolate’s cell populations at the temperature in question was delineated in terms of what percentage of cells remained metabolically active in the culture. Proportion of metabolically-active cells in a given culture was determined by testing the cells’ ability to imbibe the nonfluorescent and nontoxic molecule called fluorescein diacetate (FDA), and subsequently hydrolyze the same to the tracer compound fluorescein by means of esterase activity (89). Fluorescein-stained cells, eventually, were detected with the help of flow cytometry, as described previously (90, 91).

The procedure of fluorescence-activated cell sorting (FACS) is described below in a nutshell. Cell-pellet was harvested via cold (4°C) centrifugation for 20 minutes at 6000 *g*, and then resuspended in 2 mL of a 0.9% NaCl solution. From a 0.5% (w/v) FDA (Sigma, USA) solution in dimethyl sulfoxide, 4 µl was added to the cell suspension and incubated at 37°C for 15 minutes. The cells were then washed and resuspended again in 500 µl of 0.9% NaCl. Ultimately, with the help of a BD FACSVerse flow cytometer (Becton, Dickinson and Company, USA), 10000 cells were analyzed randomly for their ability to fluoresce through 475-495 nm excitation and 520-530 nm emission. The BD FACSuite software package (Becton, Dickinson and Company) was used to present the data in the form of a dot plot depicting the level of fluorescence recorded for each cell as a function of its photodiode-array-detected ability to forward scatter light having a wave length of 488 nm. Positions of the experiment-specific quadrant gates separating the metabolically-active cells from their inactive counterparts were determined by analyzing an unstained version of the sample.

### Testing chemoorganoheterotrophic growth on complex carbon compounds

The TMA isolates were tested for 4°C and −10°C aerobic growth on various simple to complex organic compounds as sole sources of energy, electron, and carbon. For this purpose, each strain was cultured aerobically on modified basal and mineral salts (MS) solution (92) supplemented with any one of the following organic compounds (HiMedia Laboratories, India) at one particular instance (L^−1^ double distilled Milli-Q water): acetate (10 mM), agar (2 g), benzoate (5 mM), cellulose (5 g), chitin (10 g), n-hexadecane (0.5 mL), pectin (10 g), water-soluble starch (4g) and xylan (10 g).

For each experiment, a seed culture of the test strain was prepared by as described above. From the 20 mL seed culture, cells were harvested, then washed twice with 0.9% NaCl, and eventually resuspended in 1 mL MS solution. This cell suspension in MS was added to the test medium (MS solution supplemented with any one of the 10 organic compounds mentioned above) in such a way that the specified concentration of the medium was reached only after the addition of the inoculum (final volume of the test culture was 20 mL). Experiments checking growth at 4°C were carried out in 50 mL Erlenmeyer flasks, whereas those checking growth at −10°C were carried out in 50 mL polypropylene tubes. CFU density of a culture at a given time point of incubation was determined as described for the LB-dependent growth experiments. Furthermore, each of the 27 TMA isolates was tested for its ability to grow in MS solution devoid of any organic carbon. These experiments were carried out in the same way as described above for testing growth in MS supplemented with a single organic compound,

### Testing antibiosis potential

Every TMA isolate was tested by agar overlay assay (93) for its antibiosis potentials against higher-temperature-adapted, mesophilic, Gram negative and Gram positive bacteria that had been isolated previously from warmer habitats within the Western-Himalayan and Trans-Himalayan territories. The Gram negative targets included the well-known model microorganism *Escherichia coli* K-12 (Gammaproteobacteria), plus the temperate soil isolate *Advenella kashmirensis* WT001 that belonged to Betaproteobacteria (94, 95), and the Puga Valley hydrothermal vent isolate *Paracoccus* sp. SMMA_5 that belonged to Alphaproteobacteria (86, 91). The Gram positive targets included the Chumathang hydrothermal vent isolates *Bacillus subtilis* SC_1, *Bacillus licheniformis* PAMA2_SD1, and *Lysinibacillus fusiformis* LAPE1_SD1, all of which belonged to the phylum Bacillota (Dutta et al. unpublished). Each TMA isolate was also tested for its antibiosis capabilities against other isolates obtained from the TMA.

The potential antagonists under assessment (these were taken in batches of four isolates at a time) were first grown in LB, at 15°C for 48 h, to generate their seed cultures. 10 µL inocula from the individual seed cultures were spotted on LA plates at minimum mutual distances of 3 cm, following which the plates were incubated at 15°C for 48 h. After fully grown colonies had appeared for all the potential antagonists investigated, 100 µL of a mid-log-phase culture of the target organism against which antibiosis was to be tested, was mixed with 900 µL molten agar (0.4% w/v) having 37°C temperature, and poured uniformly on the LA plates that were already dotted by incumbent colonies of the potential antagonists. Seed culture of the incoming bacterium (target of potential antibiosis) was grown at 28°C or 15°C, according as the organism was a mesophile from a foreign habitat or an isolate from the TMA. Eventually, these test plates were incubated for 48 h (again, at 28°C or 15°C, according as the incoming bacterium was a mesophile from a foreign habitat or a TMA isolate) and checked for the development of lawns of growth, or clear zones of inhibition, for the incoming organism, around the pre-established colonies of the incumbent bacteria (potential antagonists).

### Whole genome extraction, sequencing, and annotation

Whole genomic DNA was extracted from the LB-grown stationary phase culture of a given TMA isolate using HiPurA Bacterial Genomic DNA Purification Kit (Himedia, India), and sequenced using a Novaseq 6000 (Illumina Inc., USA) as well as a MinION (Oxford Nanopore Technologies, UK) platform. All the read datasets obtained in this way were submitted for public accession to the Sequence Read Archive of National Center for Biotechnology Information (NCBI), USA, under the BioProject PRJNA1335599 (BioSample accession numbers of all the depositions are given in Table S37). For every genome, 2×150 bp paired-end Illumina reads having Phred score above 20 were assembled alongside MinION reads having quality values >10, with the help of the software Unicycler v0.5.0 run in hybrid assembly mode (96). Prodigal v2.6.3 (97) was used to predict open reading frames (ORFs), or putative genes, within the assembled genome. Subsequenly, a catalog of protein-coding gene sequences (CDSs) was delineated by searching the repertoire of ORFs available, against the eggNOG database v5.0 (98), with the aid of eggNOG-mapper v2.1.9 (99), which in turn used the algorithm HMMER.

### Identification of genes concerned with the key phenotypes of the TMA isolates

To identify genes associated with low temperature adaptation, and low as well as high temperature adaptation, the eggNOG-derived CDS catalogs of the TMA isolates were searched on the basis of the information available in the literature for extreme temperature adaptation, alongside the gene orthology information curated in the Kyoto Encyclopedia of Genes and Genomes (KEGG; https://www.genome.jp/kegg/).

To identify genes encoding carbohydrate-active enzymes, the Prodigal-derived ORF catalogs of the isolates were annotated directly by searching against the CAZy and dbCAN3 databases using Diamond (100) and HMMER (101) algorithms (with default parameters for both) respectively. Subsequently, the collective findings of the two search exercises were reported as the catalogs of genes encoding carbohydrate-active enzymes.

To identify genes that are known to be central to the biosynthesis of different classes of antibiotics, each eggNOG-derived CDS catalog was searched based on the information available in the literature for antibiotic biosynthesis, plus the gene orthology information curated in the KEGG database. Furhtermore, to detect genes or gene clusters concerned with the biosynthesis of secondary metabolites, each Prodigal-derived ORF catalog was annotated using the bacterial version (v7.0) of the antibiotics and secondary metabolite analysis shell (antiSMASH) pipeline in strict detection mode (102, 103).

Antibiotic resistance genes were identified by annotating the assembled whole genome sequences of the individual TMA isolates against the Comprehensive Antibiotic Resistance Database (CARD, version 3.2.8) using the Resistance Gene Identifier (RGI, version 6.0.3) tool with its default analysis parameters (104).

### Metagenomics

From the lake-water, lake-sediment, and rock-dust samples, metagenomic DNA was extracted, and sequenced on a Novaseq 6000 using 2 × 250 bp paired-end read chemistry, as described previously (85, 105). After clipping the adapters and quality-filtering for an average Phred score ≥20, 25,000,000 read-pairs were extracted randomly from each metagenomic sequence dataset and assembled *de novo* using Megahit v1.2.9 with default parameters (106).

Within the >1000 bp contigs assembled from a given metagenome, ORFs or putative genes were annotated using Prodigal v2.6.3 in default mode (107). CAZyme-encoding genes were identified within the gene catalog obtained by searching against the CAZy and dbCAN3 databases using Diamond (100) and HMMER (101) with default parameters respectively. The collective findings of the two searches were eventually reported as the catalog of genes encoding CAZymes within the metagenome in question.

To get an idea about the habitat-wise prevalence (relative abundance) of the different TMA species for which whole genomes were sequenced, the proportion of sequence correspondence that existed between a given genome and the metagenome of the habitat under consideration was determined as follows. First, a subject (target) database was created by curating all the 15 whole genome sequences in hand; subsequently, all Q20 metagenomic reads available for a given habitat were mapped onto the subject database using Bowtie2 v2.4.5 in default mode (108). The alignment output file obtained was processed with SAMtools v 1.13 (109) to report the number of metagenomic reads that matched each genome of the target database individually.

## Supporting information

An MS Workbook containing Supplementary Tables

An MS Word file containing a Supplementary Figure and Supplementary References

## ACKNOWLEDGEMENTS

Extensive on-field assistance provided by Sri Asgar Ali of Choglamsar, Ladakh, India is gratefully acknowledged. We thank Dr. Soumya Chatterjee, Biodegradation Technology Division, Defence Research Laboratory, India, for valuable discussions on cold-customized bioreactors.

## FUNDING

The study was financed by Bose Institute through Intramural Research Grants. S.C. and M.M. received fellowships from Department of Biotechnology (DBT), Government of India (GoI). S.D. and J.S. obtained their fellowships from Council of Scientific and Industrial Research, GoI. J. G. and S. S. got fellowships from University Grants Commission, GoI. NM received fellowship from Bose Institute. Bioinformatic analyses were carried out using computational resources available under an EMR project funded by DBT, GoI (BT/PR40174/BTIS/137/45/2022).

## AUTHOR CONTRIBUTIONS

W.G. conceived the study, designed the experiments, interpreted the results, and wrote the paper. S.C. anchored the program, planned and performed the experiments, analyzed and curated the data, and also composed the paper. S.D. analyzed the data, while S.D., J.G., S.S., M.M., J.S., and N.M. performed the experiments. All authors read and approved the manuscript.

## COMPETING INTEREST

The authors declare no competing interest.

## SUPPLEMENTARY DATA

Supplementary information and data are available online in the form of a Word file named Supplimentary_Information.docx, and an Excel file named Supplimentary_Dataset.xlsx.

## DATA AVAILABILITY

GenBank accession numbers for the 16S rRNA genes of the new isolates are as follows: PV789631 (*Arthrobacter* sp. TRD_SC_1), PV789699 (*Arthrobacter* sp. TRD_SC_3), PV789726 (*Microbacterium* sp. TRD_SC_10); PV793448 (*Microbacterium* sp. TW_SC_2); PV793500 (*Microbacterium* sp. TW_SC_3); PV789771 (*Mycetocola* sp. TRD_SC_2); PV682789 (*Paenarthrobacter* sp. TRD_SC_7); PV790021 (*Pseudarthrobacter* sp. TRD_SC_9); PV793432 (*Pseudarthrobacter* sp. TS_SC_4); PV793425 (*Sanguibacter* sp. TS_SC_8); PV789776 (*Streptomyces* sp. TRD_SC_5); PV793427 (*Trichococcus* sp. TS_SC_9); PV793436 (*Flavobacterium* sp. TS_SC_2); PV793440 (*Flavobacterium* sp. TS_SC_5); PV790452 (*Ancylobacter* sp. TW_SC_1); PV793444 (*Acinetobacter* sp. TW_SC_4); PV793419 (*Aeromonas* sp. TS_SC_11); PV793374 (*Pseudomonas* sp. TS_SC_1); PV793375 (*Pseudomonas* sp. TS_SC_3); PV793377 (*Pseudomonas* sp. TS_SC_10); PV793378 (*Pseudomonas* sp. TS_SC_12); PV793393 (*Pseudomonas* sp. TS_SC_13); PV793413 (*Psychrobacter* sp. TS_SC_6). All genome and metagenome sequence data have been deposited to the NCBI under the BioProject accession number PRJNA1335599.

Sequence read datasets for the genomes have been deposited to the Sequence Read Archive (SRA), while the assembled whole genome sequences have been deposited to the GenBank, under the BioSamples accession numbers SAMN52016225 (*Acinetobacter* sp. TW_SC_4), SAMN52392035 (*Aeromonas* sp. TS_SC_11), SAMN52016223 (*Ancylobacter* sp. TW_SC_1), SAMN52016228 (*Arthrobacter* sp. TRD_SC_6), SAMN52016222 (*Cryobacterium* sp. TS_SC_7), SAMN52392036 (*Flavobacterium* sp. TS_SC_5), SAMN52016224 (*Microbacterium* sp. TW_SC_2), SAMN52016227 (*Mycetocola* sp. TRD_SC_2), SAMN52392037 (*Paenarthrobacter* sp. TRD_SC_7), SAMN52016226 (*Pseudarthrobacter* sp. TRD_SC_9), SAMN52016221 (*Pseudomonas* sp. TS_SC_3), SAMN52392038 (*Psychrobacter* sp. TS_SC_6), SAMN52016220 (*Sanguibacter* sp. TS_SC_8), SAMN52392039 (*Streptomyces* sp. TRD_SC_5), and SAMN52392040 (*Trichococcus* sp. TS_SC_9).

Sequence read datasets for the metagenomes have been deposited to the SRA under the BioSamples accession numbers SAMN53833703 (weathered rock dust), SAMN53833704 (lake-water), and SAMN53833705 (lake-sediment).

## Notes

### Competing Interest Statement

The authors have declared no competing interest.

### Summary of Updates

We have now brought about several changes in the paper to increase the robustness of the work. Specifically, we have reinforced the study with a completely fresh line of genomic evidences that were derived by sequencing de novo 15 out of the total 27 bacterial species isolated on a &primeone genome per genus&prime basis. We have also invoked the whole metagenome shotgun sequence datasets that we had already generated from the same lake-water, lake-sediment, and rock-dust samples collected on 28 October 2021, but had kept aside for a separate study of microbiome structure and function. For a ready perusal, please see the five new Result sections, the four new figures (numbered as Figs. 8, 9, 10, and 11 in the revised manuscript) and 23 new Tables (numbered as Tables 2, S37-S57, and S61 in the revised manuscript), and the corresponding new interpretations that are interspersed mainly across the the Discussion section.

