## Supplementary material for "Cryo-adapted bacterial copiotrophs from a Trans-Himalayan lake-desert ecosystem as biogeothermometers of warming, mitigators of perturbation, and candidates for biodegradation": An MS Word file containing a Supplementary Figure and Supplementary References

**Author Address:**

^$^	Faculty of Engineering and Natural Sciences, Tampere University, Finland

^#^	Department of Microbiology and Cell Science, Fort Lauderdale Research and Education Center, University of Florida, USA

**Table of Contents**

**Supplementary Figure**

**Figure S1.** Correlation between the proportions of divisible and metabolically-active cells across the species incapable of growing in Luria broth at (a) -10°C, (b) 28°C, (c) 37°C, or (d) 42°C.

**Figure S2.** Growth rates, or absence of growth, recorded for the individual TMA isolates in Luria broth at -10°C, 4°C, 15°C, 28°C, 37°C, and 42°C.

**Figure S3.** Growth temperature windows of the bacterial genera under which the different TMA isolates were classified.

**Supplementary Tables**

**(Tables S1 through S61. All SI tables are more than one page long, so they have been given as individual sheets of an Excel Workbook named Supplementary_Dataset)**

**Table S1.** Changes recorded in the CFU densities of the TMA isolates (plus *E*. *coli*) upon incubation at -10°C in Luria broth (for the cultures lacking increase in CFU density, proportions of FDA-stained cells were recorded at the end of the incubations).

**Table S2.** Changes recorded in the CFU densities of the TMA isolates (plus *E*. *coli*) upon incubation at 4°C in Luria broth.

**Table S3.** Changes recorded in the CFU densities of the TMA isolates (plus *E*. *coli*) upon incubation at 15°C in Luria broth.

**Table S4.** Changes recorded in the CFU densities of the TMA isolates upon incubation in MS-acetate (MSAc) at 4°C.

**Table S5.** Changes recorded in the CFU densities of the TMA isolates upon incubation in MS-agar (MSAg) at 4°C.

**Table S6.** Changes recorded in the CFU densities of the TMA isolates upon incubation in MS-albumin (MSAl) at 4°C.

**Table S7.** Changes recorded in the CFU densities of the TMA isolates upon incubation in MS-benzoate (MSB) at 4°C.

**Table S8.** Changes recorded in the CFU densities of the TMA isolates upon incubation in MS-cellulose (MSCl) at 4°C.

**Table S9.** Changes recorded in the CFU densities of the TMA isolates upon incubation in MS-chitin (MSCt) at 4°C.

**Table S10.** Changes recorded in the CFU densities of the TMA isolates upon incubation in MS-hexadecane (MSH) at 4°C.

**Table S11.** Changes recorded in the CFU densities of the TMA isolates upon incubation in MS-pectin (MSP) at 4°C.

**Table S12.** Changes recorded in the CFU densities of the TMA isolates upon incubation in MS-starch (MSS) at 4°C.

**Table S13.** Changes recorded in the CFU densities of the TMA isolates upon incubation in MS-xylan (MSX) at 4°C.

**Table S14.** Growth (percentage increase in CFU density) attributable to the utilization of different carbon compounds as single cheomorganoheterotrophic substrates at 4°C.

**Table S15.** Changes recorded in the CFU densities of the TMA isolates upon incubation in MS-acetate (MSAc) at -10°C.

**Table S16.** Changes recorded in the CFU densities of the TMA isolates upon incubation in MS-agar (MSAg) at -10°C.

**Table S17.** Changes recorded in the CFU densities of the TMA isolates upon incubation in MS-albumin (MSAl) at -10°C.

**Table S18.** Changes recorded in the CFU densities of the TMA isolates upon incubation in MS-benzoate (MSB) at -10°C.

**Table S19.** Changes recorded in the CFU densities of the TMA isolates upon incubation in MS-cellulose (MSCl) at -10°C.

**Table S20.** Changes recorded in the CFU densities of the TMA isolates upon incubation in MS-chitin (MSCt) at -10°C.

**Table S21.** Changes recorded in the CFU densities of the TMA isolates upon incubation in MS-hexadecane (MSH) at -10°C.

**Table S22.** Changes recorded in the CFU densities of the TMA isolates upon incubation in MS-pectin (MSP) at -10°C.

**Table S23.** Changes recorded in the CFU densities of the TMA isolates upon incubation in MS-starch (MSS) at -10°C.

**Table S24.** Changes recorded in the CFU densities of the TMA isolates upon incubation in MS-xylan (MSX) at -10°C.

**Table S25.** Growth (percentage increase in CFU density) attributable to the utilization of different carbon compounds as single cheomorganoheterotrophic substrates at -10°C.

**Table S26.** Changes recorded in the CFU densities of the TMA isolates upon incubation in minimal salts (MS) solution at 4°C.

**Table S27.** Changes recorded in the CFU densities of the TMA isolates upon incubation in minimal salts (MS) solution at -10°C.

**Table S28.** Changes recorded in the CFU densities of the TMA isolates (plus *E*. *coli*) upon incubation at 28°C in Luria broth (for the cultures lacking increase in CFU density, proportions of FDA-stained cells were recorded at the end of the incubations).

**Table S29.** Changes recorded in the CFU densities of the TMA isolates (plus *E*. *coli*) upon incubation at 37°C in Luria broth (for the cultures lacking increase in CFU density, proportions of FDA-stained cells were recorded at the end of the incubations).

**Table S30.** Changes recorded in the CFU densities of the TMA isolates (plus *E*. *coli*) upon incubation at 42°C in Luria broth (for the cultures lacking increase in CFU density, proportions of FDA-stained cells were recorded at the end of the incubations).

**Table S31.** Growth rates calculated for the individual TMA isolates at -10°C in Luria broth.

**Table S32.** Growth rates calculated for the individual TMA isolates at 4°C in Luria broth.

**Table S33.** Growth rates calculated for the individual TMA isolates at 15°C in Luria broth.

**Table S34.** Growth rates calculated for the individual TMA isolates at 28°C in Luria broth.

**Table S35.** Growth rates calculated for the individual TMA isolates at 37°C in Luria broth.

**Table S36.** Growth rates calculated for the individual TMA isolates at 42°C in Luria broth.

**Table S37.** Key attributes of the assembled whole genome sequences of the 15 TMA isolates shortlisted.

**Table S38.** eggNOG‑based annotation of all protein-coding sequences (CDSs) identified within the genome of *Arthrobacter* sp. TRD_SC_6.

**Table S39.** eggNOG‑based annotation of all protein-coding sequences (CDSs) identified within the genome of *Cryobacterium* sp. TS_SC_7.

**Table S40.** eggNOG‑based annotation of all protein-coding sequences (CDSs) identified within the genome of *Microbacterium* sp. TW_SC_2.

**Table S41.** eggNOG‑based annotation of all protein-coding sequences (CDSs) identified within the genome of *Mycetocola* sp. TRD_SC_2.

**Table S42.** eggNOG‑based annotation of all protein-coding sequences (CDSs) identified within the genome of *Paenarthrobacter* sp. TRD_SC_7.

**Table S43.** eggNOG‑based annotation of all protein-coding sequences (CDSs) identified within the genome of *Pseudarthrobacter* sp. TRD_SC_9.

**Table S44.** eggNOG‑based annotation of all protein-coding sequences (CDSs) identified within the genome of *Sanguibacter* sp. TS_SC_8.

**Table S45.** eggNOG‑based annotation of all protein-coding sequences (CDSs) identified within the genome of *Streptomyces* sp. TRD_SC_5.

**Table S46.** eggNOG‑based annotation of all protein-coding sequences (CDSs) identified within the genome of *Trichococcus* sp. TS_SC_9.

**Table S47.** eggNOG‑based annotation of all protein-coding sequences (CDSs) identified within the genome of *Flavobacterium* sp. TS_SC_5.

**Table S48.** eggNOG‑based annotation of all protein-coding sequences (CDSs) identified within the genome of *Ancylobacter* sp. TW_SC_1.

**Table S49.** eggNOG‑based annotation of all protein-coding sequences (CDSs) identified within the genome of *Acinetobacter* sp. TW_SC_4.

**Table S50.** eggNOG‑based annotation of all protein-coding sequences (CDSs) identified within the genome of *Aeromonas* *salmonicida* TS_SC_11.

**Table S51.** eggNOG‑based annotation of all protein-coding sequences (CDSs) identified within the genome of *Pseudomonas* sp. TS_SC_3.

**Table S52.** eggNOG‑based annotation of all protein-coding sequences (CDSs) identified within the genome of *Psychrobacter* sp. TS_SC_6.

**Table S53.** Number of genes identified in the shortlisted TMA isolates for adaptation to low temperatures, and low as well as high temperatures.

**Table S54.** Number of genes encompassed by the shortlisted TMA isolates under the different categories of carbohydrate-active enzymes (CAZymes).

**Table S55.** Annotation details of the key genes concerned with the biosynthesis of different antibiotics, as identified within the eggNOG based gene catalogue of the 15 TMA isolates.

**Table S56.** AntiSMASH-based annotation details of the genes concerned with secondary metabolites biosynthesis, as identified within the genomes of the 15 TMA isolates.

**Table S57.** CARD-RGI-based annotation details of the genes concerned with antibiotic resistance, as identified within the genomes of the 15 TMA isolates.

**Table S58.** Members retrieved previously from other cold/frigid habitats of the world for the genera that were currently isolated from the Tso Moriri area.

**Table S59.** Members retrieved previously from discrete mesic and hot habitats for the genera that were currently isolated from the Tso Moriri area.

**Table S60.** Minimum and maximum growth temperatures known (in the literature) thus far for the global strains of the genera that were isolated in this study from the Tso Moriri area.

**Table S61.** Proportion of metagenomic reads from the three distinct habitats of the Tso Moriri lake-desert ecosystem that matched with sequences from the genomes of the 15 representative isolates (randomly extracted 50,000,000 reads were tested from each metagenome for their alignment with the genome sequences).

**Table S62.** Number of genes detected under the different categories of carbohydrate-active enzymes, within the assembled metagenomes of Tso Moriri water, Tso Moriri sediment, and weathered rock dust covering the lake-side hill.

**Supplementary References**

**References used in Table S58**

**References used in Table S59**

**References used in Table S60**

**Supplementary Figure**

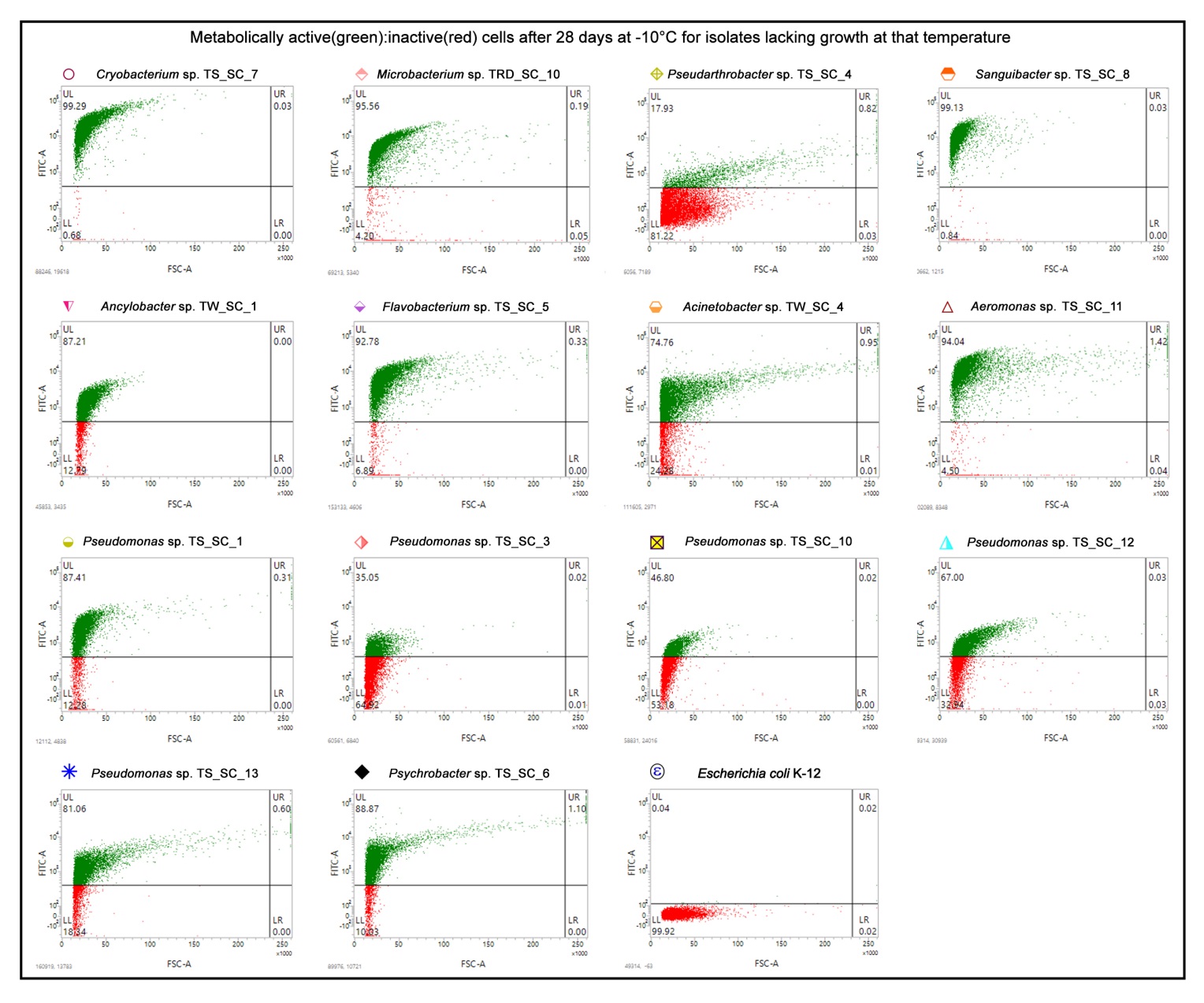

**Fig. S1.** Flow cytometry-derived dot plots depicting what proportions of cells were stained by FDA (after 28 days of incubation) in the LB cultures of the 14 TMA isolates incapable of growing at -10°C. In every dot plot, the levels of fluorescence recorded for 10000 randomly chosen cells have been shown as the functions of the corresponding forward scattering data, with green and red points representing FDA stained and unstained cells respectively. Each FACS experiment was repeated two more times, with deviation from the result (percentages of FDA stained and unstained cells) shown here being <2% on every occasion.

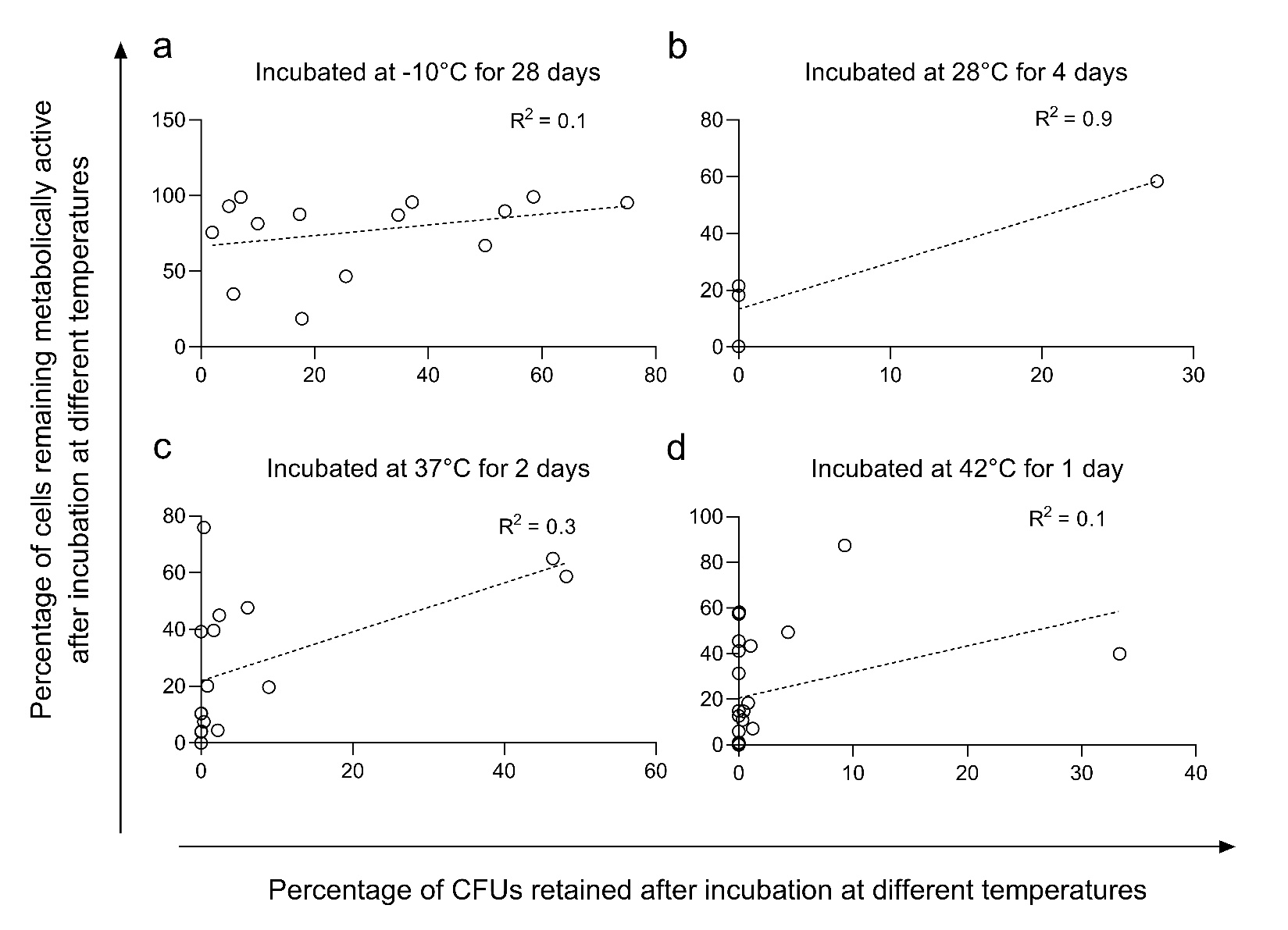

**Fig. S2.** Correlation between the proportions of divisible and metabolically-active cells across the species incapable of growing in Luria broth at (**a**) -10°C, (**b**) 28°C, (**c**) 37°C, or (**d**) 42°C.

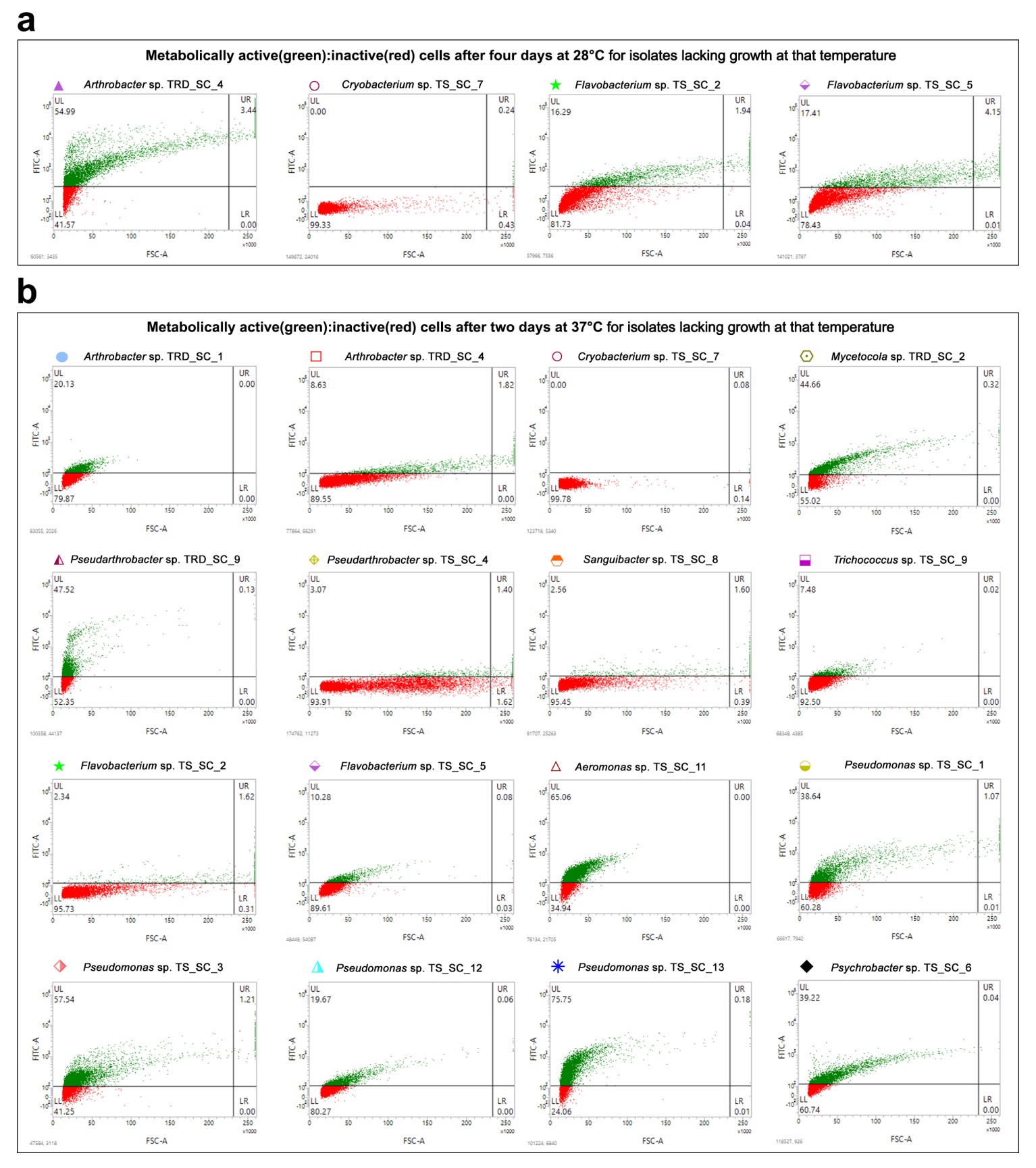

**Fig. S3.** Flow cytometry-derived dot plots depicting what proportions of cells were stained by FDA in the LB cultures of the isolates incapable of growing after (**a**) four days of incubation at 28°C, and (**b**) two days of incubation at 37°C. In each dot plot, the fluorescence levels recorded for 10000 randomly chosen cells have been shown as the functions of the corresponding forward scattering data, with green and red points representing FDA stained and unstained cells respectively. Each FACS experiment was repeated two more times, with deviation from the result (percentages of FDA stained and unstained cells) shown here being <2% on every occasion.

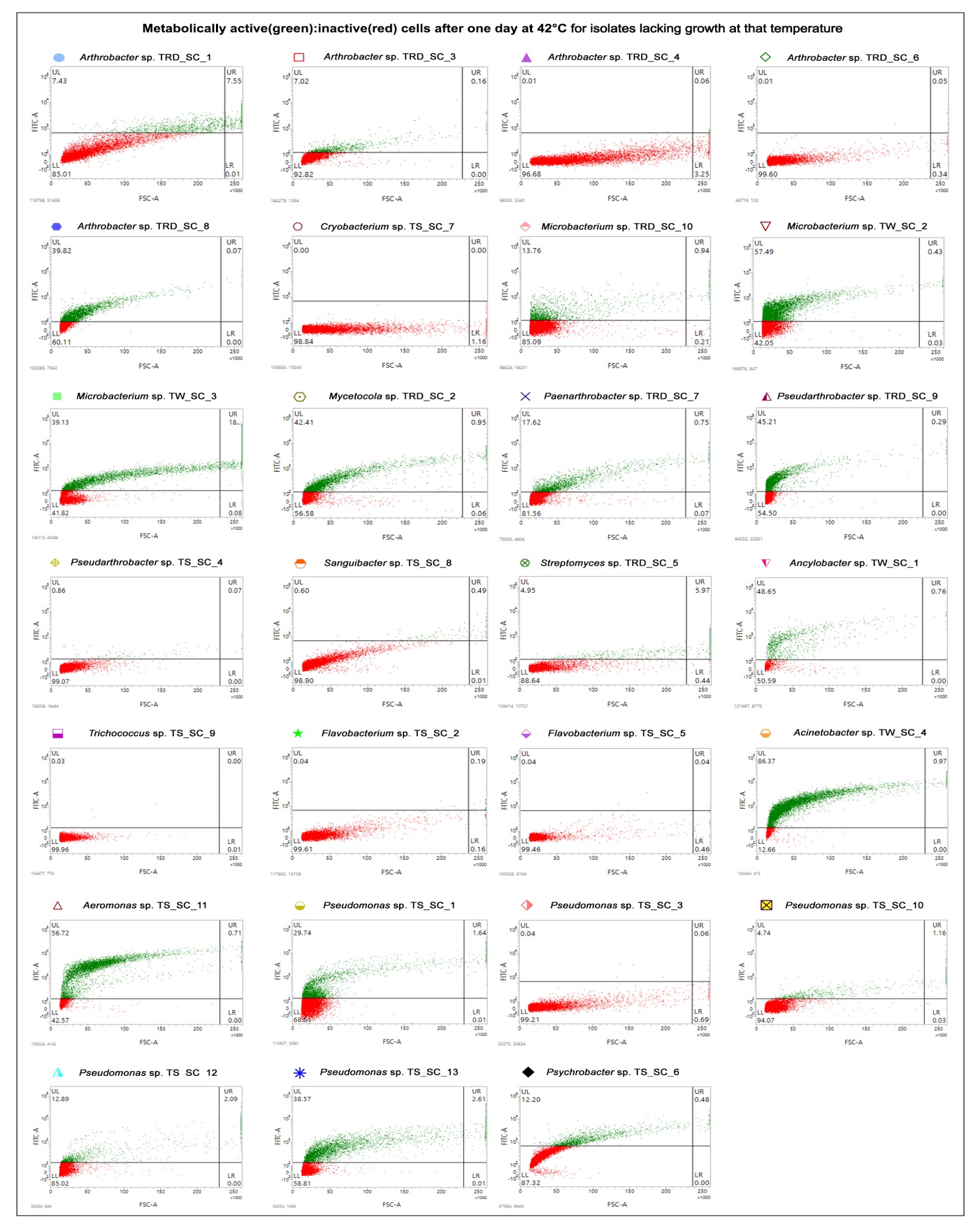

**Fig. S4.** Flow cytometry-derived dot plots depicting what proportions of cells were stained by FDA in the LB cultures of the 27 TMA isolates after 24 h incubation at 42°C**.** In each plot, the fluorescence levels recorded for 10000 randomly chosen cells have been shown as the functions of the corresponding forward scattering data, with green and red points representing FDA stained and unstained cells respectively. Each FACS experiment was repeated two more times, with deviation from the result (percentages of FDA stained and unstained cells) shown here being <2% on every occasion.

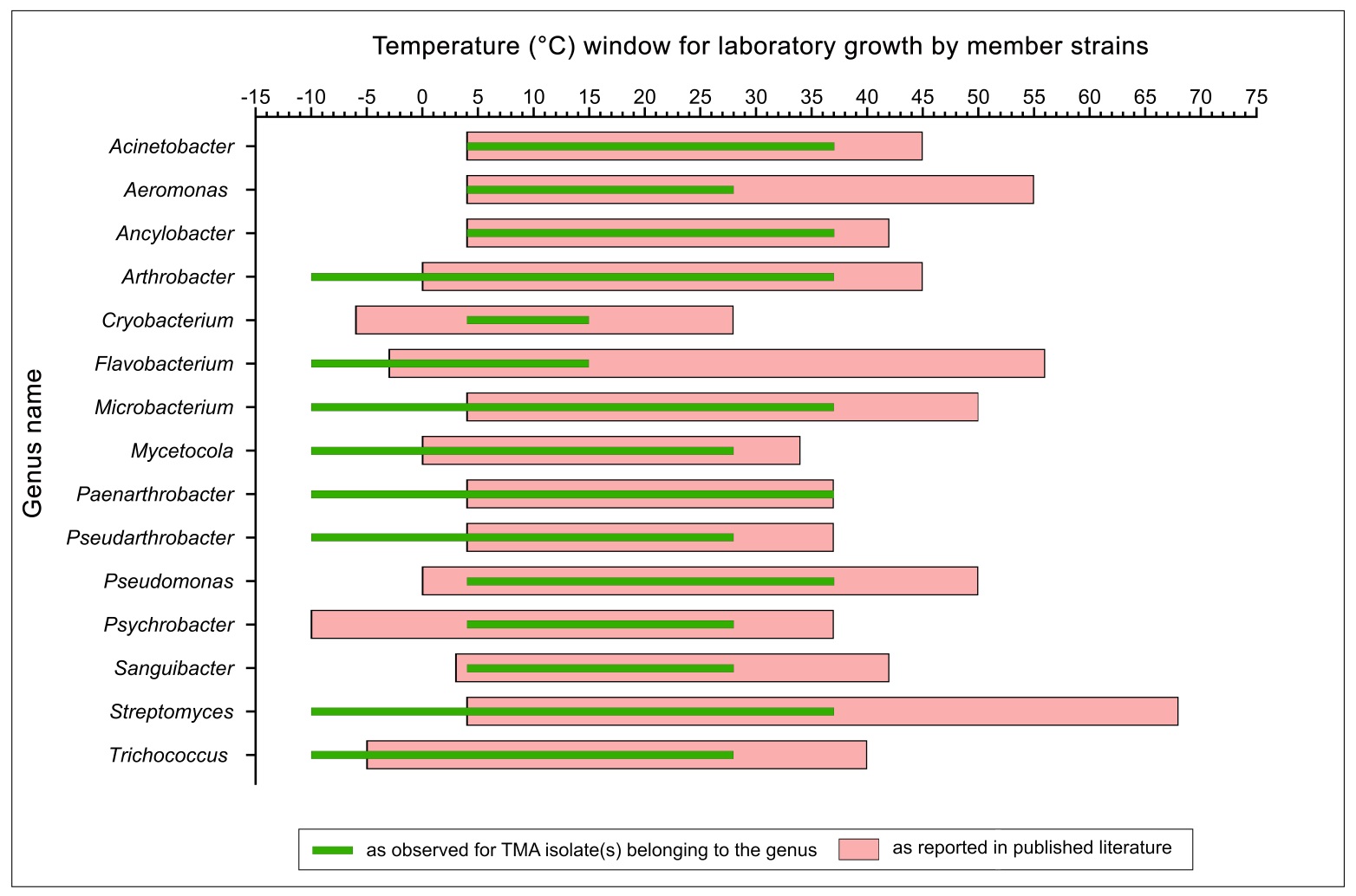

**Fig. S5.** Growth temperature windows of the bacterial genera under which the different TMA isolates were classified. Data obtained for the present isolates have been depicted in the context of the data available in the literature for their global relatives (see Table S39, and Supplementary References, for the detailed information).
